# Enhanced PIEZO1 Function Contributes to the Pathogenesis of Sickle Cell Disease

**DOI:** 10.1101/2025.03.19.643952

**Authors:** Luis O. Romero, Manisha Bade, Laila Elsherif, Jada D. Williams, Xiangmei Kong, Adebowale Adebiyi, Kenneth I. Ataga, Shang Ma, Julio Cordero-Morales, Valeria Vásquez

## Abstract

Sickle cell disease (SCD), an inherited blood disorder caused by a mutation in the β-globin gene, is characterized by sickle erythrocytes that are prone to hemolysis, causing anemia and vaso-occlusion crises. In sickle erythrocytes, hemoglobin aggregation is followed by altered cation permeability and subsequent dehydration. Interventions to restore cation permeability can decrease hemolysis and ameliorate the symptoms associated with SCD. PIEZO1 is a non-selective mechanosensitive cation channel that regulates erythrocyte volume. Gain-of-function (GOF) mutations in PIEZO1 cause hemolytic anemia by increasing cation permeability, leading to erythrocyte dehydration in humans and mice. Although PIEZO1 plays a key role in erythrocyte homeostasis, its role in SCD remains unknown. Here, we demonstrate that the function of the PIEZO1 channel is upregulated in sickle erythrocytes of humans and mice, and this enhancement can be restored through a dietary intervention. We found that PIEZO1 function in sickle erythrocytes resembles that of the GOF mutation causing hemolytic anemia. A diet enriched in the *ω*-3 fatty acid eicosapentaenoic (EPA) acid decreases PIEZO1 function in sickle erythrocytes and hemolysis in a mouse model of SCD. Furthermore, EPA decreases hemolysis and reduces inflammatory markers. We propose that PIEZO1 contributes to the increase in nonselective cationic conductance (i.e., Psickle), which leads to dehydration downstream of hemoglobin polymerization. Our results suggest that reducing PIEZO1 function is a promising therapeutic approach to reestablishing normal cation permeability in SCD.

## Introduction

Sickle cell disease (SCD) is an inherited condition characterized by a mutation in the gene that codes for the hemoglobin subunit β (i.e., sickle hemoglobin), causing erythrocytes to adopt a sickle-like shape. ^1^ This lifelong and debilitating illness disproportionately impacts Black or African American populations, with the Centers for Disease Control and Prevention reporting an incidence of 1 in 365 births within this demographic. ^2^ Hematopoietic stem cell transplantation, and potentially gene therapy and gene editing approaches, offer curative treatments for SCD. ^3^ Beyond the technical complications (e.g., well-matched donors, age, cost), these options are inaccessible to most patients. Current therapeutic strategies aim to alleviate symptoms and modify the severity of the disease by decreasing hemolytic anemia and vaso-occlusive complications with approved drug therapies (hydroxyurea, L-glutamine, and Crizanlizumab) as well as red blood cell transfusions. ^1,3^ Despite the availability of newer therapeutic options, the vast majority of individuals with SCD, especially those residing in sub-Saharan Africa, have very limited access to these treatments. ^4^

Hemoglobin polymerization results in the formation of rigid, fiber-like structures that underlie the pathogenesis of SCD. ^5,6^ Sickling results in less deformable and more adhesive erythrocytes, resulting in hemolytic anemia, hypoxemia, red blood cell dehydration, and blockage of blood flow. These can cause vaso-occlusive pain crises, organ damage, and endothelial dysfunction. ^1,4,7,8^ Biomarkers of constant hemolysis include increased plasma hemoglobin and indirect bilirubin levels, as well as the presence of inflammatory markers (e.g., cytokines and chemokines). ^9–11^ The probability of sickle hemoglobin polymerizing is directly proportional to a decrease in the intracellular oxygen concentration and exponentially proportional to the intracellular sickle hemoglobin concentration. ^12^ Interestingly, hemoglobin polymerization alters erythrocyte membrane organization (e.g., loss of membrane asymmetry) and ion permeability. ^1,13–15^ Several membrane proteins are dysfunctional in this context (e.g., K^+^-Cl^-^ cotransporters, Na^+^-K^+^ pump, ion channels) ^16,17^ and are linked to a nonspecific cationic conductance, activated downstream of deoxygenation, termed Psickle. ^18–21^ Psickle leads to dehydration and a subsequent rise in intracellular hemoglobin concentration. This pathophysiological shift is critical in the erythrocyte sickling process, which leads to hemolysis. Identifying the ion channels contributing to Psickle is crucial for developing new therapeutic strategies.

PIEZO1 is a mechanosensitive, non-selective cation channel that regulates erythrocyte volume. ^22–25^ Lack of PIEZO1 leads to overhydrated erythrocytes and eliminates mechanically induced calcium influx, as demonstrated by calcium imaging experiments. ^24^ Whereas gain-of-function (GOF) point mutations in the *PIEZO1* gene underlie human hereditary xerocytosis, in which the mutant channel increases cation permeability, leading to dehydrated erythrocytes. ^22,25–28^ A mouse model of a *Piezo1* GOF mutation causing dehydrated xerocytosis mirrors several pathological features observed in SCD, including reduced osmotic fragility and anemia. ^25^ Sickle erythrocytes exhibit reduced intracellular K^+^ and elevated intracellular Ca^2+^.^29,30^ Mechanical stress activates PIEZO1 channels, causing an increase in intracellular Ca^2+^ levels, which in turn opens the Ca^2+^-activated potassium channel (i.e., Gárdos channel). ^24^ Whether PIEZO1 is a potential target for regulating Ca^2+^ homeostasis in sickle erythrocytes remains to be determined.

Patients with SCD have abnormal erythrocyte membranes characterized by elevated ω-6 fatty acid levels and reduced ω-3 fatty acids. ^31–34^ We have previously established that PIEZO1 function is modulated by the lipid composition of the plasma membrane and that it can be fine-tuned by enriching specific dietary fatty acids *in vitro*. ^35^ Additionally, we have shown that PIEZO2 — a homolog of PIEZO1 — can be modulated *in vivo* by diets enriched in polyunsaturated fatty acids, alleviating symptoms in two unrelated neurogenetic disorders. ^36,37^ Given the common features of dehydration and increased cation permeability associated with SCD and PIEZO1 GOF mutations, as well as the fatty acid imbalance in SCD blood cell membranes, we aim to determine whether PIEZO1 channel function is altered in SCD. Here, we demonstrate that PIEZO1 function is enhanced in sickle erythrocytes from human adults and mice with SCD in a sex-independent manner. Notably, we restored normal PIEZO1 function by feeding a mouse model of SCD with a diet enriched in eicosapentaenoic acid (EPA). This dietary intervention decreased plasma hemoglobin and indirect bilirubin levels and reduced various inflammatory markers. Our findings suggest that PIEZO1 is a key contributor to the altered cation permeability observed in SCD and emphasize the therapeutic potential of targeting this channel to alleviate symptoms associated with SCD and other hematological disorders.

## Materials and Methods

### Ethics approval

The participation of human donors was approved by the Institutional Review Board of the University of Tennessee Health Science Center (UTHSC; IRB 20-07604-XP). All participants provided written informed consent. Mice procedures described below were reviewed and approved by the UTHSC Institutional Animal Care and Use Committee (UTHSC IACUC number: Protocol #19-0084 and #22-0320) and the University of Texas Health Science Center (UT Houston; AWC-23-0093). All methods were carried out following approved guidelines.

### Human samples

Patients between ages 18 to 65, with confirmed diagnoses of sickle cell anemia (11 HbSS and one HbSβ^0^ thalassemia), who had no severe pain episodes requiring medical contact during the preceding 4 weeks donated a blood sample while in their non-crisis, steady states. Subjects were recruited during routine clinic visits at the Methodist Comprehensive Sickle Cell Center in Memphis, Tennessee. Participants were excluded if they were pregnant, on anticoagulant therapy, or if less than 3 months had passed since their last transfusion. Blood was collected on sterile BD vacutainer blood collection tubes with K2 EDTA and used for experiments within 4 h of collection.

### Mice

Male and female Townes transgenic HbSS (hα/hα::β^S^/β^S^; sickling) mice were obtained from the Jackson laboratory or bred in-house from heterozygous breeders (hα/hα::β^A^/β^S^; coming from strain #013071). ^38^ C57BL/6 J mice (WT) were obtained from The Jackson Laboratory (Stock No. 000664). *Piezo1* GOF (R2482H; in the hematopoietic system) mice were generated by breeding *Piezo1*^cx/cx^ ^25^ with *Vav1-cre* (The Jackson Laboratory, stock# 018968). GOF *Piezo1* (R2482H) mice were generated and maintained on the C57BL/6 background (these animals were backcrossed at least 10 generations to C57BL/6). Littermates were used as control animals for experiments involving GOF *Piezo1* mice. Adult (2- to 6-month-old) mice were housed with a 12 h light/dark cycle with food and water *ad libitum*.

Blood from mice was obtained through a tail vein or cardiac puncture. Blood collected through the tail vein puncture was diluted in the solution used for electrophysiological experiments and used within 4 h of collection. Blood collected through cardiac puncture was collected in K2 EDTA tubes and used within 4 h of collection. For GOF *Piezo1* or their WT littermates drawn blood was collected, in UT Southwestern Medical Center at Dallas, into tubes containing acid-citrate-dextrose buffer (Sigma-Aldrich C3821), centrifuged at 100 *g* for 15 min at room temperature, supernatant, and buffy coat were discarded and the erythrocyte pellet was resuspended and washed with PBS two-three times. After the final wash, erythrocytes were diluted in 0.9% NaCl saline, kept on ice packs, and shipped overnight to UT Houston. These erythrocytes were used for electrophysiological recordings within 18 h of collection.

### Electrophysiology

Patch-clamp recordings were performed in the inside-out configuration in the voltage-clamp mode. The bath solution contained 140 mM KCl, 6 mM NaCl, 2 mM CaCl_2_, 1 mM MgCl_2_, 10 mM glucose, and 10 mM HEPES (pH 7.4), while the pipette solution contained 140 mM NaCl, 6 mM KCl, 2 mM CaCl_2_, 1 mM MgCl_2_, 10 mM glucose, and 10 mM HEPES (pH 7.4). Pipettes were made of borosilicate glass (Sutter Instruments) and fire-polished to a resistance between 8 and 10 MΩ before use. Mechanical stimulation was performed using the voltage-clamp (constant −60 mV, unless otherwise noted). Recordings were sampled at 100 kHz and low pass filtered at 2 kHz using a MultiClamp 700 B amplifier and Clampex (Molecular Devices, LLC). Yoda1 (Tocris Bioscience), and GsMTx-4 (Abcam) stocks were prepared using DMSO and water, respectively, and diluted in the pipette solution to face the membrane from the outer leaflet. Leak currents before mechanical stimulation were subtracted offline from the current traces with ClampFit (Molecular Devices, LLC). Recordings with leak currents >50 pA, and patches with giga-seals that did not withstand at least five consecutive steps of mechanical stimulation were excluded from analyses.

### Mechanical stimulation

For pressure-clamp assays, excised membrane patches were mechanically stimulated with negative pressure applied through the patch pipette using a High-Speed Pressure Clamp (ALA Scientific), which was automated using a MultiClamp 700B amplifier through Clampex. Inside-out patches were probed using a square-pulse protocol consisting of a 50 ms 10-mmHg pre-pulse, immediately followed by incremental pressure steps (−10 mmHg), each lasting 200 ms in 1-s intervals. For current-voltage (IV) relationships, a square-pulse protocol consisting of a 1-s -120 mmHg pressure step was applied every 10 seconds at a voltage ranging from −60 to 60 mV.

### Diet supplementation

HbSS (6-8 weeks old) mice were pair-fed for 16 weeks with a high-fat diet (anhydrous milk fat supplemented, Dyets # 1050181), modified AIN-93G purified rodent diet with 59% fat-derived calories from anhydrous milk fat (kcal/kg): casein (716), L-cystine (12), maltose dextrin (502), cornstarch (818.76), anhydrous milk fat (2430), soybean oil (630), mineral mix (# 210025; 30.8), and vitamin mix (#310025; 38.7) or an ω-3 (enriched in EPA; Dyets Inc. #112246) modified AIN 93G enriched diet with 59% fat derived calories from menhaden oil (kcal/kg): casein (716), L-cystine (12), maltose dextrin (502), cornstarch (818.76), menhaden oil (2430), soybean oil (630), mineral mix (#210025; 30.8), and vitamin mix (#310025; 38.7).

### Mice blood collection

Blood from mice fed with high fat or ω-3 diet was collected in a micro-sample tube EDTA K3E (Sarstedt Inc, Cat#41.1504.005) by cardiac puncture. For plasma collection, the blood was centrifuged at 2,000 *g* for 15 min at 4°C. The plasma was separated and stored at -80°C. The buffy coat was discarded. The bottom-packed erythrocytes were collected and stored at -80°C. For serum collection, mouse blood was collected in a micro-sample tube serum CAT (Sarstedt Inc, Cat #41.1392.105). The serum micro-sample tube was left in an upright position undisturbed for 20 min at room temperature allowing the blood to clot. Then, the blood was centrifuged at 1,500 *g* for 15 min at 4 °C. Serum was separated, transferred into new microcentrifuge tubes, stored at -80°C, and thawed at the time of assay.

### Liquid chromatography-mass spectrometry

The bottom-packed erythrocytes collected from mice blood fed with high fat or ω-3 diet were shipped to Wayne State University. Total and free fatty acids were quantified at the Lipidomics Core Facility at Wayne State University.

### Plasma Hemoglobin

Mice plasma hemoglobin concentration was measured with the QuantichromTM Hemoglobin Assay Kit (BioAssay systems, Cat #DIHB-250) following the manufacturer’s instructions.

### Serum chemistry analyses

Mice serum chemistry parameters were obtained using a Beckman Coulter AU480 Automated analyzer at the Center of Comparative Medicine at Baylor College of Medicine.

### Mouse cytokine profile assay

Serum cytokine profiles were determined with the Mouse Cytokine Array Panel kit (R&D Systems, Cat #ARY006). 100 μL of serum sample was used for each array. The serum sample was diluted in array buffers and mixed with reconstituted mouse cytokine array Panel A detection antibody cocktail. The sample antibody mixture was incubated for 1 h at room temperature and was added to the nitrocellulose membrane. The membrane was incubated overnight at 4°C on a rocking platform shaker. Streptavidin-HRP was diluted (1:2,000) in array buffer added to the membrane after three washes and incubated for 30 min at room temperature. Chemiluminescent reagents mix was added to the membrane and then imaged in a ChemiDoc Touch Imaging System (Bio-Rad). The mean pixel density of membrane spots representing each cytokine were analyzed using Image Lab Software (v6.1.0 build 7; Bio-Rad).

### Plasma haptoglobin

Plasma samples were thawed at the time of assay and processed according to the haptoglobin ELISA kit (Aviva Systems biology, Cat #OKIA00095). The plasma samples were diluted and treated in microtiter wells coated with anti-haptoglobin (anti-HPT) antibodies. Anti-HPT antibodies conjugated with horseradish peroxidase were added after washing. Then, a chromogenic substrate solution was added. Samples absorbance was measured at 450 nm with an Infinite M200 PRO NanoQuant microplate reader (Tecan).

### Hematological indices

Whole blood samples were processed within 30 min of collection on a VETSCAN® HM5 Hematology Analyzer (Zoetis) as per the manufacturer’s instructions to obtain the following parameters: erythrocyte count, hematocrit, total hemoglobin (i.e., plasma plus intracellular), mean corpuscular hemoglobin, and mean corpuscular volume.

### Data analysis, statistics, and reproducibility

Data were plotted using OriginPro (2018 v:b9.51.195; OriginLab Corp.). The time constant of deactivation τ was obtained by fitting a single exponential function (1):

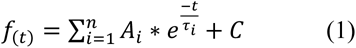

where A = amplitude, τ = time constant, and the constant y-offset C for each component i. All boxplots show mean (square), median (bisecting line), bounds of the box (75^th^ to 25^th^ percentiles), and outlier range with 1.5 coefficient (whiskers). Statistical analyses were performed using GraphPad InStat software (version 3.10; GraphPad Software Inc.). We used the Kolmogorov and Smirnov method to determine data distribution, as well as Bartlett’s test to determine differences between standard deviations. Individual tests are described in each of the Figure legends. No statistical method was used to predetermine the sample size. No data were excluded from the analyses. The experiments were not randomized. The investigators were blind to genotype and treatment whenever possible. Experiments were performed at least three times on different days from different/independent preparations.

### Data sharing

The source data underlying the figures and supplementary figures will be made available as a Source Data file on Figshare upon acceptance of the article for publication.

## Results

### Humans with sickle cell disease display increased PIEZO1 currents

To measure PIEZO1 activity in humans with SCD, we collected blood from 12 adult donors during routine clinic visits and patch-clamped their erythrocytes in the inside-out configuration. Individuals with SCD have a varying range (4-44%) of sickle erythrocytes in their blood. ^39,40^ This feature allowed us to measure PIEZO1 activity in sickle and non-sickle erythrocytes under isogenic conditions (**Figure 1A** and **supplemental Figure 1A**). For all SCD donors, regardless of the sex, PIEZO1 mechanocurrents were larger in sickle erythrocytes than in non-sickle ones at every pressure pulse (**Figures 1A-B** and **supplemental Figure 1B-E**). Our results demonstrate that PIEZO1 function is increased in sickle erythrocytes.

**Figure 1.**
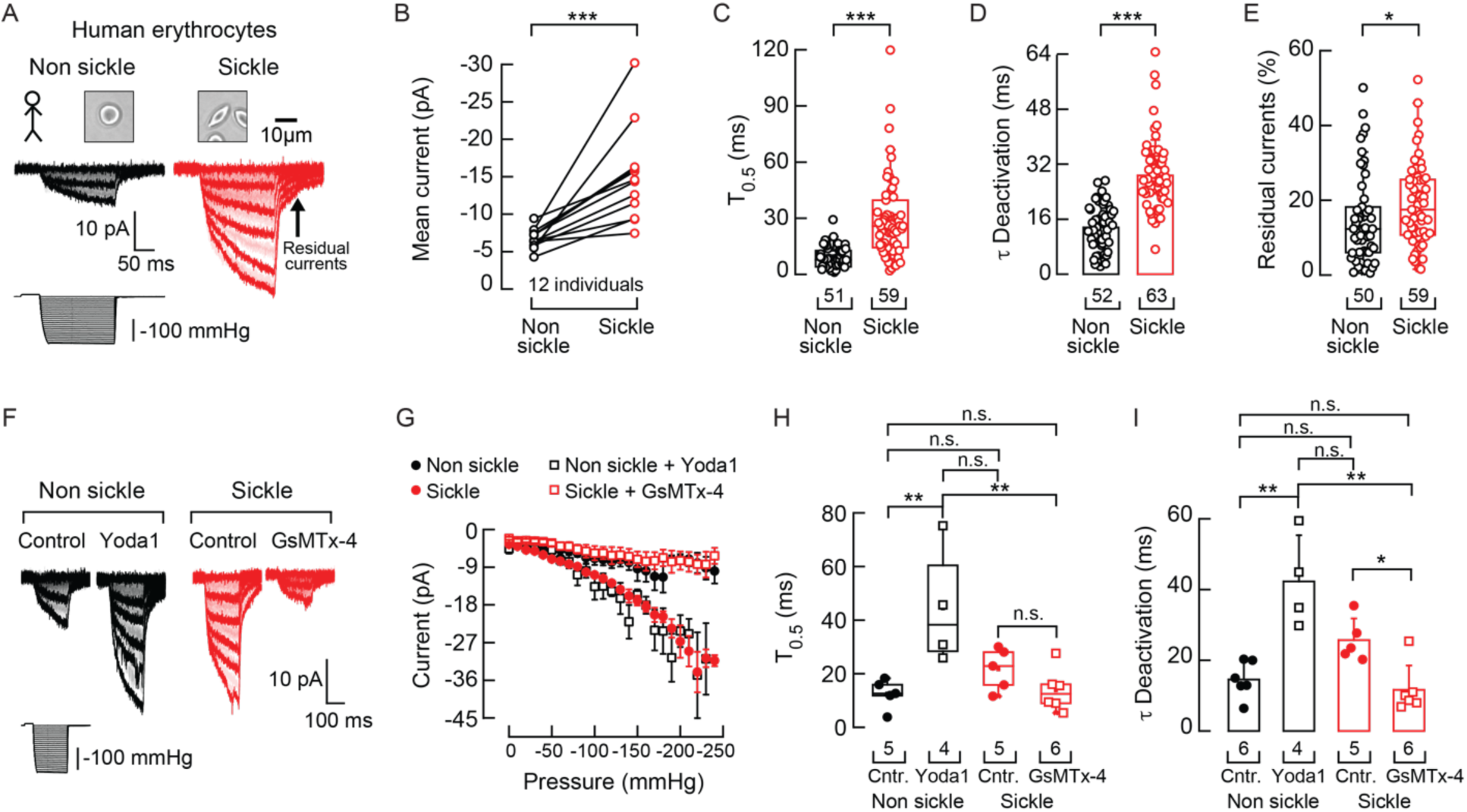
Increased mechano-activated currents in sickle erythrocytes from SCD patients. **A**. Top: Representative bright field micrographs of sickle and non-sickle erythrocytes from a human with SCD. Bottom: Representative inside-out recordings elicited by negative pressure square pulses at a constant voltage of −60 mV. **B**. Mean mechanocurrents elicited by -130 mmHg. Currents are paired per individual. Two-tailed Wilcoxon matched-pairs signed-ranks test (*p* = 0.0005). **C.** Time required to reach half of the mechanocurrents maximal value (T_0.5_) elicited by -130 mmHg. Two-tailed Mann-Whitney test (U = 410, *p* = 2.71^-12^). **D**. Time constant of deactivation (τ) elicited by maximum negative pressure. Two-tailed Unpaired t-test (*t* = 8.97, *p* = 7.43^-15^). **E**. Percentage of peak current 30 ms after the -130-mmHg stimulus ends. Two-tailed Mann-Whitney test (U = 1100, *p* = 0.023). **F**. Representative inside-out recording elicited by negative pressure square pulses at a constant voltage of −60 mV of control (DMSO) or Yoda1 (30 µM)-exposed non-sickle and control or GsMTx-4 (7 µM)-exposed sickle erythrocytes. **G**. Current-pressure relationships (elicited at -60 mV) of control (DMSO) or Yoda1-exposed non-sickle and control or GsMTx-4 exposed sickle erythrocytes. Symbols are mean ± SEM. **H**. Time required to reach half of the mechanocurrents maximal value (T_0.5_) elicited by -130 mmHg from control (DMSO) or Yoda1-exposed non-sickle and control or GsMTx-4 exposed sickle erythrocytes. Kruskal-Wallis (H = 9.79, *p* = 0.02) with Dunn’s multiple comparison test. **I**. Time constant of deactivation (τ) elicited by the maximum negative pressure from control (DMSO) or Yoda1-exposed non-sickle and control or GsMTx-4 exposed sickle erythrocytes. Kruskal-Wallis (H = 14.06, *p* = 0.0028) with Dunn’s multiple comparison test. Bars are mean ± SD. Boxplots show the mean, median, and 75^th^ to 25^th^ percentiles. n is denoted above the *x*-axis. Asterisks indicate values significantly different from the control (∗p < 0.05, ∗∗p < 0.01, and ∗∗∗p < 0.001) and n.s. indicates not significantly different.

We further quantified PIEZO1 gating behavior to identify the salient functional differences between sickle and non-sickle erythrocytes. In sickle erythrocytes, PIEZO1 takes longer to reach half-maximal activation and deactivation (**Figure 1C-D**). We also observed higher residual currents after the mechanical stimuli end (**Figure 1A, E and supplemental Figure 1F**). We did not detect differences in the latency of response — the time between the onset of mechanical stimulation and the initiation of PIEZO1 mechanocurrents — between sickle and non-sickle erythrocytes (**supplemental Figure 1G**). Likewise, the linear current-voltage relationships and reversal potentials in both erythrocyte types are similar (**supplemental Figure 2A-B**). These results indicate that the primary changes in the PIEZO1 function in sickle erythrocytes occur in current amplitudes, deactivation, and residual currents.

We leveraged the available pharmacology of the PIEZO1 channel to further characterize its function in human erythrocytes. Specifically, we challenged membranes of non-sickle erythrocytes with Yoda1, a selective positive modulator of PIEZO1 activity, ^41^ while treating sickle erythrocytes with GsMTx-4, a nonspecific inhibitor of mechanosensitive ion channels. ^42^ Importantly, Yoda1 elicited large mechanocurrents in non-sickle erythrocytes, comparable to those observed in sickle erythrocytes at every pressure step (**Figure 1F-G and supplemental Figure 2C**). Yoda1 also slowed the time to reach half-maximal activation and deactivation and increased the residual currents, similar to those measured in control (DMSO) sickle erythrocytes (**Figure 1H-I and supplemental Figure 2D-E**). On the other hand, GsMTx-4 diminished mechanocurrents and half-maximal activation while also accelerating deactivation to levels comparable to those observed in non-sickled cells (**Figure 1F-I**). Neither compound affected the latency of current onset to mechanical stimulation (**supplemental Figure 2F**). Taken together, our functional and pharmacological characterization demonstrates that the PIEZO1 channel mediates an increased cationic permeability in human sickle erythrocytes.

### PIEZO1 function in sickle erythrocytes resembles a PIEZO1 GOF mutation

The functional features of PIEZO1 in human and mouse sickle erythrocytes are reminiscent of the human PIEZO1 GOF mutations that cause xerocytosis, previously characterized when transfected in mammalian cell lines. ^43^ To directly measure the currents of PIEZO1 GOF in erythrocytes, we performed patch-clamp experiments from a hematopoietic lineage-specific mouse line expressing the GOF *Piezo1 R2482H* ^25^ (**Figure 2A**) and its corresponding littermates. Following mechanical stimulation, the GOF channel exhibited larger currents, slower activation kinetics, and residual currents than WT (**Figure 2B-F**). On the other hand, we did not find differences in the τ of deactivation (**Figure 2G**). The enhanced mechanosensitivity and persistent currents observed in erythrocytes expressing the *R2482H* mutation resemble the functional features we measured for PIEZO1 in human sickle erythrocytes (**Figure 1**). Parenthetically, we carried out a pharmacological characterization of PIEZO1 currents in mouse erythrocytes. Consistent with our results in humans, we found that Yoda1 increased PIEZO1 mechanocurrents, and GsMTx-4 reduced the currents in wild-type (WT) mouse erythrocytes (**supplemental Figure 3**). Together, our results represent the first macroscopic characterization of PIEZO1 mechanocurrents in human and mouse erythrocytes.

**Figure 2.**
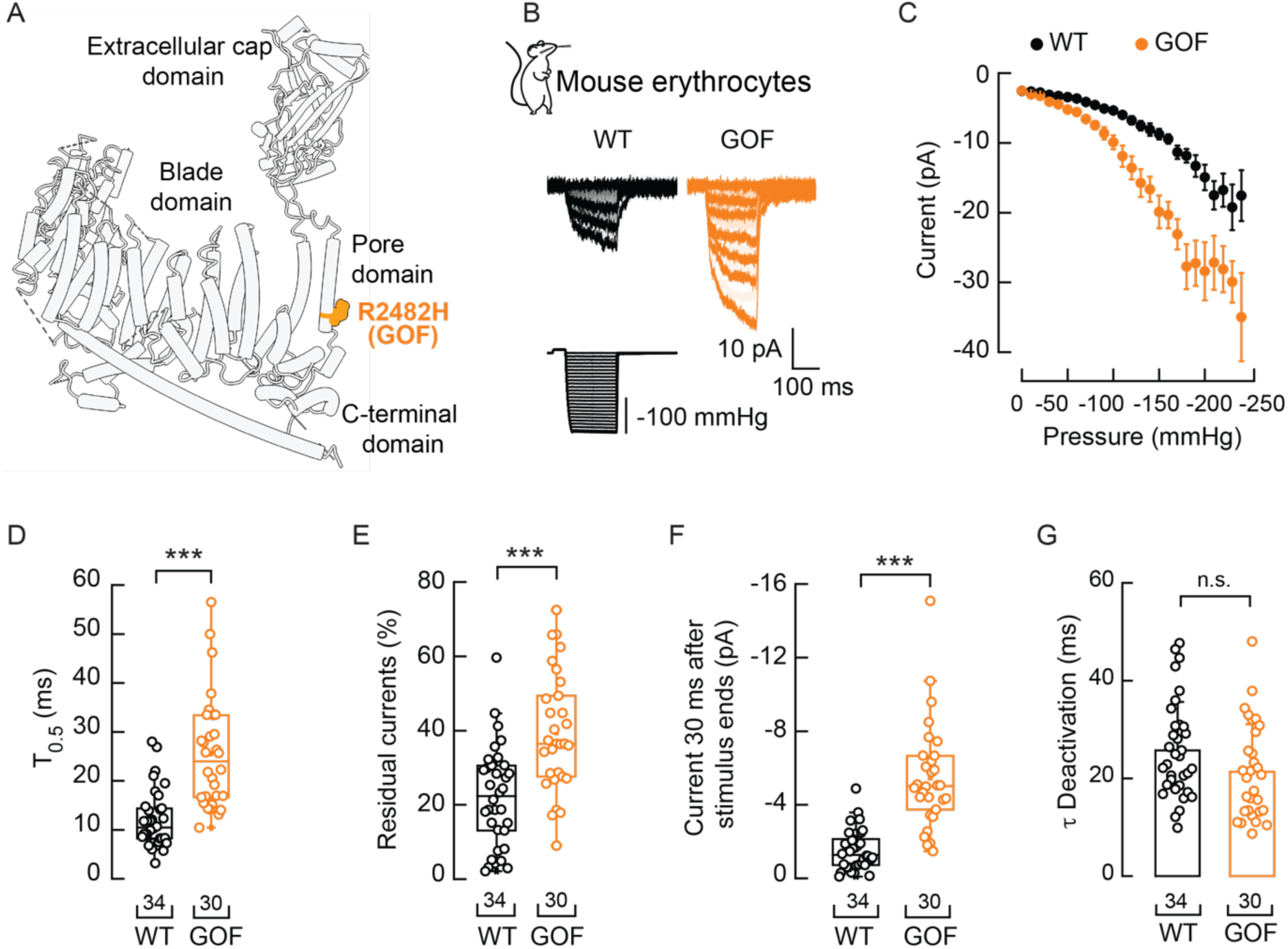
Electrophysiological characterization of PIEZO1 GOF mutation R2482H in mouse erythrocytes. **A.** Cylinder representation of mouse PIEZO1 monomer (PDB ID: 5Z10) highlighting an equivalent residue that when mutated (R2482H; GOF) in human PIEZO1 causes xerocytosis. **B.** Representative inside-out recordings elicited by negative pressure square pulses at a constant voltage of −60 mV from wild-type (WT) and PIEZO1 GOF mice erythrocytes. **C.** Current-pressure relationships (elicited at -60 mV) from WT (n = 34) and PIEZO1 GOF (n = 30) mice erythrocytes. Symbols are mean ± SEM. **D.** Time required to reach half of the mechanocurrents maximal value (T_0.5_) elicited by -130 mmHg. Two-tailed Unpaired *t*-test (*t* = 120, *p* = 1.53^-8^). **E.** % of peak current 30 ms after the -130-mmHg stimulus ends. Two-tailed Unpaired *t*-test (t = 4.7, p = 1.48^-5^). **F.** Current left 30 ms after the -130-mmHg stimulus ends. Two-tailed Unpaired *t*-test (t = 6.22, p = 4.77^-8^). **G**. Time constant of deactivation (τ) elicited by maximum negative pressure. Two-tailed Unpaired t-test (*t* = 1.78, *p* = 0.08). Boxplots show the mean, median, and 75^th^ to 25^th^ percentiles. n is denoted above the *x*-axis. Asterisks indicate values significantly different from the control (∗∗∗*p* < 0.001). n.s. indicates not significantly different.

### Elevated PIEZO1 currents in a mouse model of sickle cell disease

Our studies on SCD erythrocytes in humans provided valuable insights into the underlying pathophysiology of this disorder. Next, we shifted our focus to a mouse model of SCD to determine potential strategies for reducing PIEZO1 function *in vivo*. We measured mechanocurrents on freshly isolated blood samples from young adult Townes mice (i.e., a mouse model of SCD; HbSS, hα/hα::β^S^/β^S^). Our results show larger PIEZO1 mechanocurrents in sickle erythrocytes compared to non-sickled cells in a sex-independent manner (**Figure 3A-D and supplemental Figure 4A-B**). Likewise, mouse PIEZO1 in sickle erythrocytes takes longer to reach half-maximal activation and deactivation, as well as higher residual currents (**Figure 3D-G and supplemental Figure 4C**). Similar to humans, we did not find differences in the latency of responses or current-voltage relationships of PIEZO1 in sickle and non-sickle mouse erythrocytes (**supplemental Figure 4D-F**). Overall, our results demonstrate that the Townes mouse model replicates the increased PIEZO1 activity observed in sickle erythrocytes of humans with SCD.

**Figure 3.**
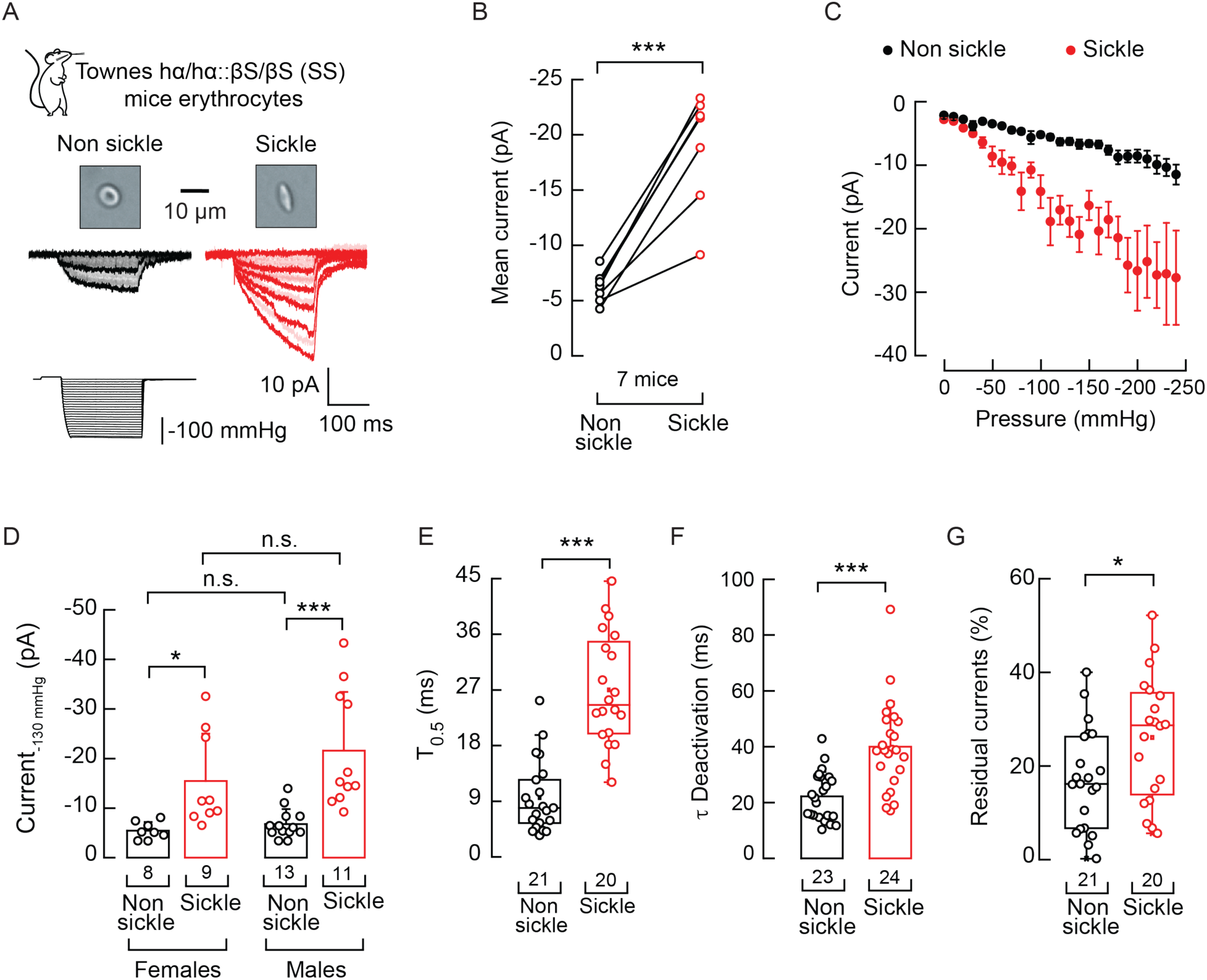
Enhanced PIEZO1 function in sickle erythrocytes from Townes SCD mice. **A**. Top: Representative bright field micrographs of non-sickle and sickle erythrocytes from the Townes mouse model of sickle cell disease. Bottom: Representative inside-out recordings elicited by negative pressure square pulses at a constant voltage of −60 mV from non-sickle and sickle erythrocytes. **B**. Mean mechanocurrents elicited by -130-mmHg. Currents are paired per mouse. Paired *t*-test (*t* = 7.44, *p* = 0.0005). **C**. Current-pressure relationships (elicited at -60 mV) from non-sickle (n = 24) and sickle (n = 23) erythrocytes, from 7 mice. Symbols are mean ± SEM. **D.** Mechanocurrents elicited by - 130 mmHg from erythrocytes from male and female mice. Two-way ANOVA with Sidakholm multiple comparison test (F = 2.23; *p* = 0.14). **E**. Time required to reach half of the mechanocurrents maximal value (T_0.5_) elicited by -130 mmHg. Two-tailed Mann-Whitney-test (U = 20, *p* = 2.02^-8^). **F**. Time constant of deactivation (τ) elicited by the maximum negative pressure. Two-tailed Mann-Whitney-test (U = 79, *p* = 2.89^-5^). **G.** % of peak current 30 ms after the -130-mmHg stimulus ends. Two-tailed Unpaired *t*-test (*t* = 2.35, *p* = 0.024). Bars are mean ± SD. Boxplots show the mean, median, and 75^th^ to 25^th^ percentiles. n is denoted above the *x*-axis. Asterisks indicate values significantly different from the control (∗∗∗*p* < 0.001 and ∗*p* < 0.05) and n.s. indicates not significant.

### An Eicosapentaenoic Acid-Enriched Diet Restores PIEZO1 Function in Sickle Erythrocytes

We have previously shown that dietary fatty acids can modulate the mechanical response of PIEZO channels *in vitro*, *ex vivo*, and *in vivo*. ^35–37,44^ Specifically, eicosapentaenoic acid (EPA) decreases PIEZO1 activity by accelerating channel inactivation. ^35^ Given the elevated PIEZO1 function observed in sickle erythrocytes of the Townes mouse model, we hypothesized that an EPA-enriched diet could restore normal PIEZO1 function. We pair-fed Townes mice for 16 weeks with a menhaden oil-enriched diet (high in EPA) or a control isocaloric high-fat diet (high in anhydrous milk fat and soybean oil). EPA accumulates in mouse tissues when its consumption in the diet is increased. ^36,45^ As expected, Townes mice fed the EPA-enriched diet exhibited higher EPA membrane content than those fed the control diet (**Figure 4A-B** and **Table 1**). Remarkably, the EPA-enriched diet reduced PIEZO1 mechanocurrents in sickle erythrocytes to levels similar to those in non-sickled cells (**Figure 4C-D**). Furthermore, the EPA-enriched diet restored PIEZO1 half-maximal activation and deactivation and reduced residual currents to levels equivalent to non-sickle erythrocytes (**Figure 4E-F and supplemental Figure 5A-B**). Neither dietary intervention affected the latency of PIEZO1 response to mechanical stimulation (**supplemental Figure 5C**). These results demonstrate that an EPA-enriched diet can effectively restore normal PIEZO1 function in sickle cells.

**Figure 4.**
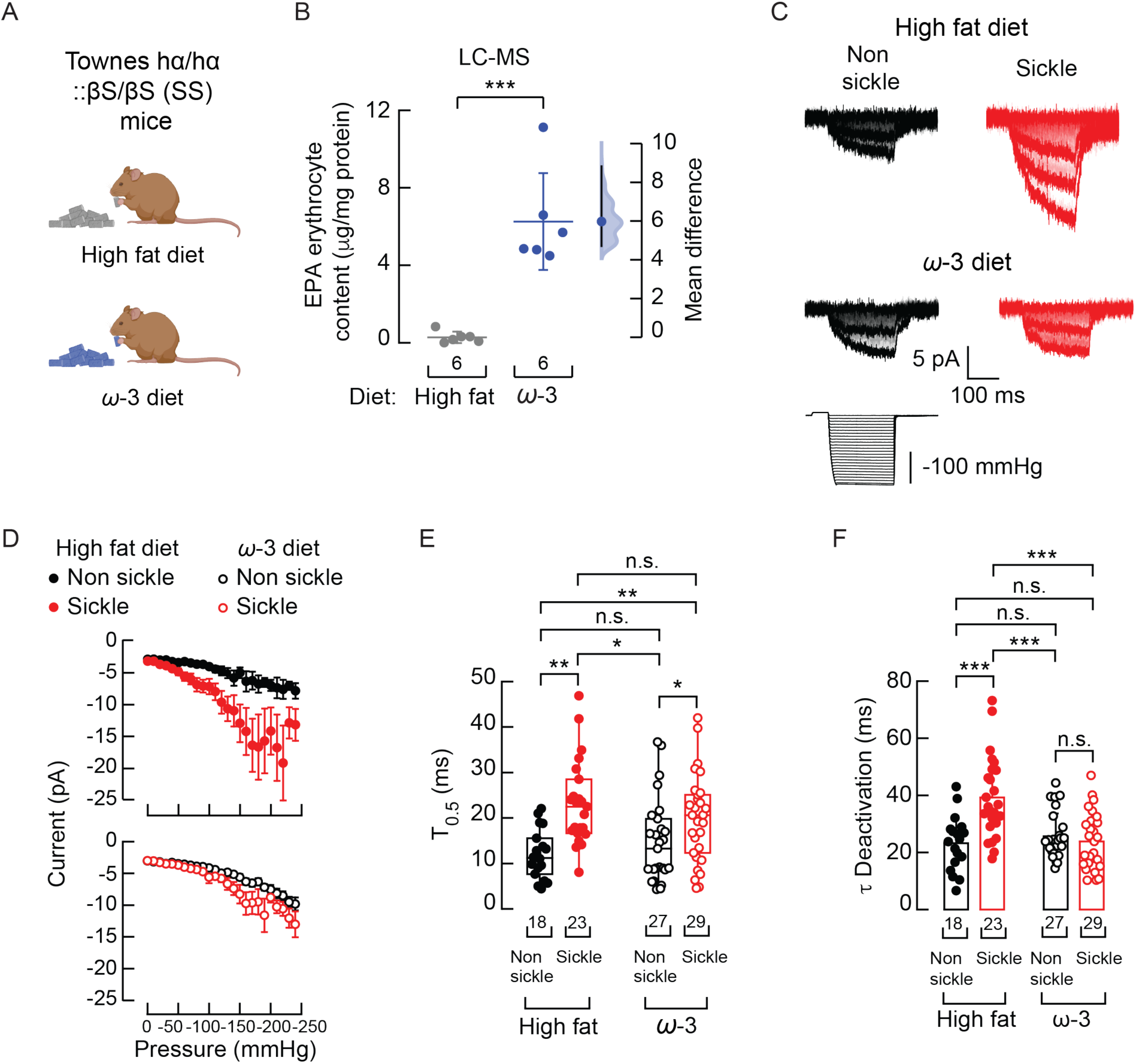
An EPA-enriched diet reduces PIEZO1 currents in sickle erythrocytes. **A**. Mouse cartoons representing the diets were created with BioRender.com. **B.** Cumming estimation plot showing the mean difference in the EPA membrane content of erythrocytes from the Townes mouse model of sickle cell disease fed with isocaloric high-fat or ω-3 enriched diets, as determined by LC-MS. Two-tailed Unpaired *t*-test (*t* = 5.81, *p* = 0.0002). **C.** Representative inside-out recordings elicited by negative pressure square pulses at a constant voltage of −60 mV. **D**. Top: Current-pressure relationships (elicited at -60 mV) from non-sickle (n = 18) and sickle (n = 23) erythrocyte from mice fed with a high-fat diet from 10 mice Bottom: Current-pressure relationships (elicited at -60 mV) from non-sickle (n = 27) and sickle (n = 29) erythrocytes from mice fed with an ω-3 enriched diet, from 8 mice. Symbols are mean ± SEM. **E**. Time required to reach half of the mechanocurrents maximal value (T_0.5_) elicited by -130 mmHg. Two-way ANOVA with Tukey multiple comparison test (F = 20.38; *p* = 1.86^-5^). **F**. Time constant of deactivation (τ) elicited by the maximum negative pressure. Two-way ANOVA with Tukey multiple comparison test (F = 8.23; *p* = 0.0051). Bars are mean ± SD. Boxplots show the mean, median, and 75^th^ to 25^th^ percentiles. n is denoted above the *x*-axis. Asterisks indicate values significantly different from the control (∗p < 0.05, ∗∗p < 0.01 and ∗∗∗p < 0.001) and n.s. indicates not significantly different.

### An EPA-Enriched Diet Decreases Hemolysis and Reduces Inflammatory Markers in a SCD Mouse Model

Based on the relationship between PIEZO1 enhanced function and hemolysis, ^25^ we sought to determine whether decreasing PIEZO1 activity with an EPA-enriched diet could improve hematological parameters associated with SCD pathophysiology. Notably, mice fed the EPA-enriched diet exhibited lower plasma hemoglobin (**Figure 5A**). Concomitant with the decrease in plasma hemoglobin, we also observed a reduction in indirect bilirubin concentration (as released hemoglobin is converted into bilirubin; **Figure 5B**). Both results support the notion that restoring PIEZO1’s normal function reduces sickle cell hemolysis. On the other hand, we did not observe differences in parameters such as erythrocyte count, hematocrit, total hemoglobin (i.e., plasma plus intracellular), and mean corpuscular hemoglobin (**supplemental Figure 6A-J**). Our results support that an EPA-enriched diet reduces PIEZO1-mediated hemolysis in SCD.

**Figure 5.**
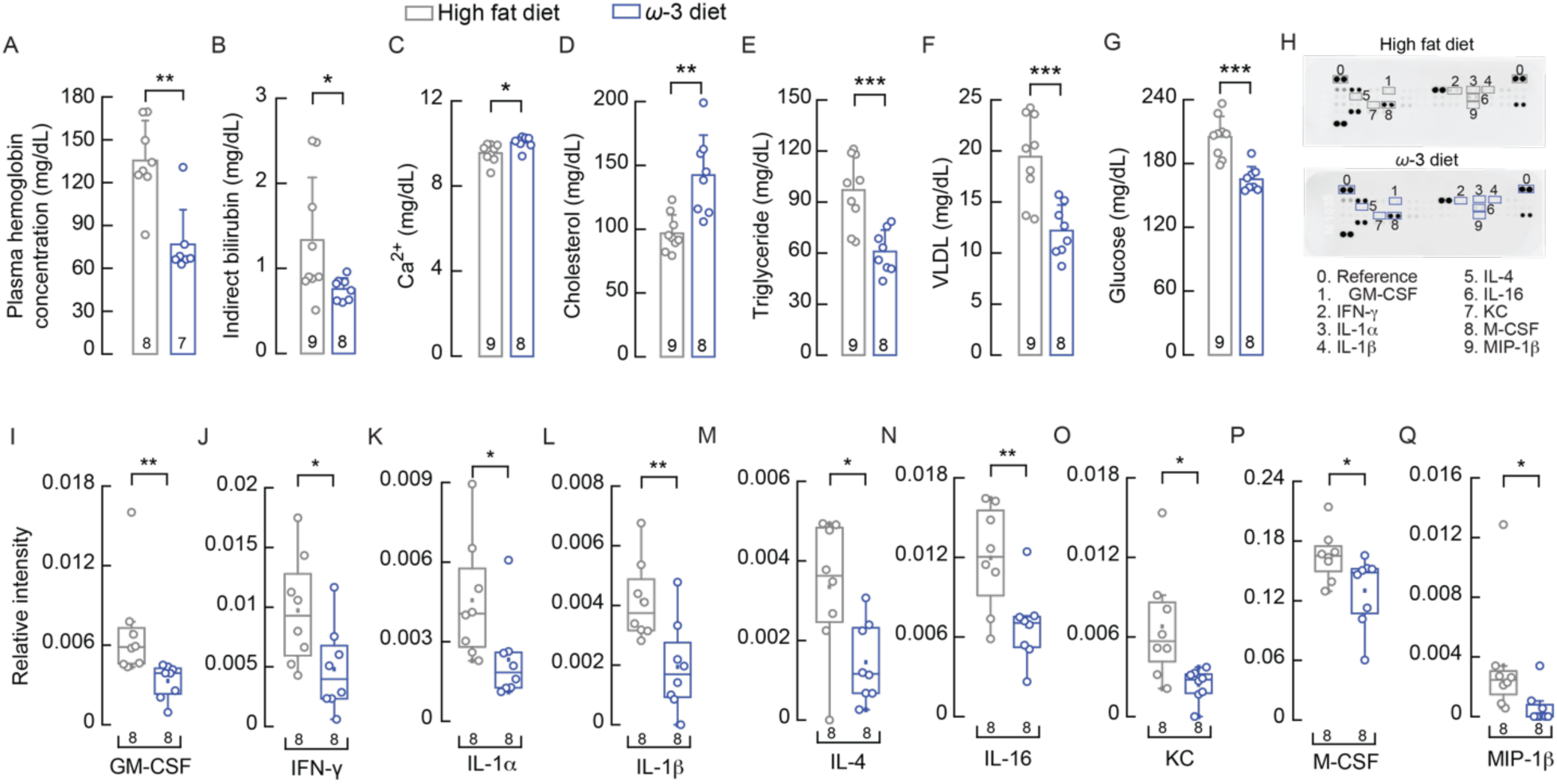
A dietary intervention reduces hemolysis and inflammation. **A.** Plasma hemoglobin concentration from mice fed with isocaloric high-fat or ω-3 enriched diets. Two-tailed Mann-Whitney test (U = 52, *p* = 0.0037). **B.** Serum indirect bilirubin concentration. Two-tailed Mann-Whitney test (U = 12.5, *p* =0.0267). **C.** Serum calcium concentration. Two-tailed Unpaired *t*-test (*t* = 2.622, *p* = 0.0192). **D.** Serum cholesterol concentration. Two-tailed Unpaired *t*-test (*t* = 3.939, *p* = 0.0013). **E.** Serum triglyceride concentration. Two-tailed Unpaired *t*-test (*t* = 4.263, *p* = 0.0007). **F.** Serum very low-density lipoprotein (VLDL) concentration. Two-tailed Unpaired *t*-test (*t* = 4.261, *p* = 0.0007). **G.** Serum glucose concentration Two-tailed Unpaired *t*-test (*t* = 5.0784, *p* = 0.0001). **H.** Representative blood serum cytokine profiles and their quantification from SS mice fed with isocaloric high-fat or ω-3 enriched diets. **I.** Relative intensity of serum granulocyte-macrophage colony-stimulating factor (GM-CSF). Two-tailed Mann-Whitney test (U = 1.5, *p* = 0.0016). **J.** Relative intensity of serum interferon-gamma (IFN-γ). Two-tailed Unpaired *t*-test (*t* = 2.390, *p* = 0.0315). **K.** Relative intensity of serum interleukin-1α (IL-1α). Two-tailed Mann-Whitney test (U = 9, *p* = 0.0148). **L.** Relative intensity of serum interleukin-1β (IL-1β). Two-tailed Unpaired *t*-test (*t* = 3.07, *p* = 0.0083). **M.** Relative intensity of serum interleukin-4 (IL-4). Two-tailed Unpaired *t*-test (*t* = 2.734, *p* = 0.0161). **N.** Relative intensity of serum interleukin-16 (IL-16). Two-tailed Unpaired *t*-test (*t* = 3.011, *p* = 0.0093). **O.** Relative intensity of serum keratinocyte-derived chemokine CXCL1(KC). Two-tailed Unpaired *t*-test (*t* = 2.87, *p* = 0.0123). **P.** Relative intensity of serum macrophage colony-stimulating factor (M-CSF). Two-tailed Mann-Whitney test (U = 12, *p* = 0.0379). **Q.** Relative intensity of serum macrophage inflammatory protein-1 beta (MIP-1β). Two-tailed Mann-Whitney test (U = 9, *p* = 0.0165). Bars are mean ± SD. Boxplots show the mean, median, and 75^th^ to 25^th^ percentiles. Asterisks indicate values significantly different from the control (∗*p* < 0.05, ∗∗*p* < 0.01. and ∗∗∗*p* < 0.001) and n.s. indicates not significantly different.

Since an EPA-enriched diet lowers free hemoglobin (which contributes to oxidative stress and inflammation), we aimed to determine its potential beneficial effects on inflammatory and metabolic markers. To this end, we conducted various analyses, including serum chemistry and cytokine profiling. We found that serum calcium and cholesterol levels rose with the EPA-enriched diet (**Figure 5C-D**). Importantly, SCD patients often exhibit hypocholesterolemia, ^46^ while hypocalcemia^47^ occurs less frequently. Additionally, triglycerides, very low-density lipoprotein (VLDL), and glucose decreased with the EPA-enriched diet (**Figure 5E-G**) —an important finding given that these parameters are typically elevated in SCD patients. ^46^ We also report other parameters that showed no significant changes (e.g., potassium, albumin, lactate dehydrogenase; **supplemental Figure 6K-S**). Notably, inflammatory markers contributing to chronic inflammation, endothelial dysfunction, and vaso-occlusive pathology were significantly reduced following the dietary intervention. These included granulocyte-macrophage colony-stimulating factor (GM-CSF), interferon-gamma (IFN-γ), interleukin-1α (IL-1α), interleukin-1β (IL-1β), interleukin-4 (IL-4), interleukin-16 (IL-16), keratinocyte-derived chemokine CXCL1 (KC), macrophage colony-stimulating factor (M-CSF), and macrophage inflammatory protein-1 β (MIP-1β; **Figure 5H-Q**). Other inflammatory markers decreased after the dietary intervention, albeit not significantly (**Supplemental Figure 7**). Overall, the dietary intervention provides multiple benefits by reducing hemolysis, stabilizing serum and lipid chemistry, and lowering inflammatory markers.

## Discussion

Our study provides insights into the role of PIEZO1 in SCD, highlighting the therapeutic potential of targeting this mechanosensitive ion channel to alleviate some of the symptoms associated with this disease. We demonstrated that PIEZO1 currents are enhanced in both humans and a humanized mouse model with SCD in a sex-independent manner. This functional upregulation mirrors the GOF mutation in PIEZO1 that causes hemolytic anemia, suggesting a common pathological mechanism involving increased cation permeability and erythrocyte dehydration.

Multiple pieces of indirect evidence suggest that PIEZO1 might play a role in SCD. For instance, 1) Cahalan et al. showed that PIEZO1 transduces mechanical forces to regulate mouse erythrocyte volume and calcium concentration, and erythrocytes lacking PIEZO1 fail to permeate calcium in response to mechanical stimuli; ^24^ 2) Vaisey et al. recorded stretch-dependent single-channel currents in a mouse erythrocyte with a conductance matching that of PIEZO1 in transfected cells; ^48^ 3) GOF mutations in PIEZO1 cause hereditary hemolytic anemia in both humans and mouse models; ^22,25–28^ and 4) Chemical sensitization of PIEZO1 with Yoda1 increases the sickling propensity of human erythrocytes. ^49^ However, no direct evidence showed whether PIEZO1 function is altered in SCD. Our electrophysiological characterization of mechanically activated currents in erythrocytes from humans and mice with SCD directly demonstrates that the function of PIEZO1 is increased in this context. This enhancement likely contributes to the nonspecific cationic conductance known as Psickle, which is activated downstream of hemoglobin polymerization. ^1^ Our study shows that inhibiting PIEZO1 function could decrease the abnormal calcium influx that enhances the Gárdos channel activity, leading to erythrocyte dehydration. Therefore, by targeting PIEZO1, we could potentially disrupt this pathological pathway and improve the clinical outcomes for SCD patients.

We now consider which factors in sickle erythrocytes increase PIEZO1 function. When erythrocytes undergo sickling, numerous changes could provoke this enhancement in channel function. Erythrocytes experience significant macroscopic shape changes upon sickling, transitioning from their normal biconcave disc shape to elongated and rigid forms. ^1^ Since PIEZO1 activity and conformation can be modulated by global curvature ^50–52^, this dramatic change in morphology could expose PIEZO1 to membrane curvatures that favor its opening. The erythrocyte’s membrane also undergoes significant mesoscopic alterations. Membrane erythrocytes from patients with SCD are characterized by unbalanced phospholipid remodeling pathways (i.e., Lands’ cycle), ^53^ abnormal membrane fatty acid composition, ^31^ loss of phospholipid asymmetry (i.e., exposure of phosphatidylserine), ^54^ and activation of sphingomyelinase and enrichment of its downstream product ceramide. ^55^ Many of these factors are known to increase PIEZO1 channel function in other cell types. For instance, *ω*-6 polyunsaturated fatty acids enrichment enhances PIEZO1 function in human microvascular endothelial cells, ^35^ surface flip-flop of phosphatidylserine activates PIEZO1 in human myotubes, ^56^ and increased sphingomyelinase activity and ceramide content enhance the prolonged activation of PIEZO1 in mouse mesenteric endothelial cells. ^57^ Our study lays the foundation for further examination of these pathways to define the mechanisms that enhance PIEZO1 function in SCD, potentially leading to novel treatments.

One of our key findings is that EPA significantly reduces the function of PIEZO1 in sickle erythrocytes. This reduction leads to decreased hemolysis (evidenced by lower plasma hemoglobin and indirect bilirubin levels) and reduced inflammatory markers. The significant decrease of inflammatory markers, including GM-CSF, IFN-γ, IL-1α, IL-1β, IL-4, IL-16, KC, M-CSF, and MIP-1β, underscores the anti-inflammatory effect of EPA. Chronic inflammation is a hallmark of SCD, contributing to endothelial dysfunction and vaso-occlusive crises. ^1,4^ By lowering these inflammatory markers, EPA may help alleviate some of the debilitating symptoms of SCD. It is still unclear whether these broader physiological benefits are only due to the inhibition of PIEZO1 function or if they also result from EPA pleiotropic effects on the vascular system and its anti-inflammatory properties. ^45,58,59^

A clinical trial of the Gárdos channel inhibitor Senicapoc in SCD patients reduced hemolysis but was terminated early due to insufficient clinical benefits. ^60,61^ Other clinical trials showed that an *ω*-3 dietary intervention, consisting of more docosahexaenoic acid (DHA) than EPA, reduced the frequency of painful vaso-occlusive crises. ^62–65^ However, the clinical and biochemical evidence remains inconclusive to date. ^64^ Our study in mice identified EPA (but not DHA, Table 1) as a key player in downregulating PIEZO1 function. This distinction is important because we and others have demonstrated that DHA has the opposite effect on PIEZO1 function by decreasing inactivation (i.e., increasing function) compared to EPA. ^35,66^ We propose that increasing EPA content in erythrocytes, alongside existing treatments for SCD, could improve the effectiveness of clinical interventions. Future research should focus on clinical trials to evaluate the efficacy of a highly purified form of EPA recently approved by the FDA (i.e., icosapent ethyl or Vascepa) ^67–69^ in SCD patients, as well as investigate other potential inhibitors of PIEZO1 function.

## Acknowledgments

This work is dedicated to the memory of Dr. Jesús G. Romero M., whose pioneering work in red blood cell electrophysiology inspired our study. We thank Dr. M. Idowu for experimental advice and Dr. T.W. Mills for access to the Tecan Infinite M200 PRO NanoQuant microplate reader. We thank Drs. V. Jayaraman and G. Wu for critically reading the manuscript. We thank members of the Vásquez and Cordero laboratories for technical support. We utilized the lipidomics Core Facility at Wayne State University (NIH S10RR027926) and The Research Services Laboratory in the Center of Comparative Medicine at Baylor College of Medicine. This work was supported by the National Institutes of Health (R01HL151735 to A.A., R35GM149218 to J.F.C.-M. and R35GM153208 to V.V.).

## Authorship Contributions

V.V is the lead author. L.O.R., J.F.C.-M, and V.V. designed the research and wrote the original draft. L.O.R., M.B., L.E., and V.V. designed the methodology. L.O.R. and M.B. performed the research, collected the data, and performed statistical analysis. L.O.R., M.B, and V.V. analyzed and interpreted data. L.E., J.D.W, X.K, A.A, K.I.A, and S.M. contributed vital new reagents. L.O.R, M.B., L.E., J.D.W, X.K, A.A, K.I.A J.F.C.-M, and V.V. review and edited the manuscript.

## References

1. Kato GJ, Piel FB, Reid CD, et al. Sickle cell disease. Nat Rev Dis Primers. 2018;4:18010.

2. Prevention. CfDCa. Data and Statistics on Sickle Cell Disease.; Retrieved August 12, 2024.

3. National Heart L, and Blood Institute. Sickle Cell Disease: Treatment; Retrieved August 12, 2024.

4. Gladwin MT, Kato GJ, Novelli EM. Sickle Cell Disease. New York, NY: McGraw-Hill Education; 2021.

5. Pauling L, Itano HA, et al. Sickle cell anemia a molecular disease. Science. 1949;110(2865):543–548.

6. Eaton WA. Linus Pauling and sickle cell disease. Biophys Chem. 2003;100(1-3):109–116.

7. Jang T, Poplawska M, Cimpeanu E, Mo G, Dutta D, Lim SH. Vaso-occlusive crisis in sickle cell disease: a vicious cycle of secondary events. J Transl Med. 2021;19(1):397.

8. Manwani D, Frenette PS. Vaso-occlusion in sickle cell disease: pathophysiology and novel targeted therapies. Blood. 2013;122(24):3892–3898.

9. Francis RB, Jr., Haywood LJ. Elevated immunoreactive tumor necrosis factor and interleukin-1 in sickle cell disease. J Natl Med Assoc. 1992;84(7):611–615.

10. Keikhaei B, Mohseni AR, Norouzirad R, et al. Altered levels of pro-inflammatory cytokines in sickle cell disease patients during vaso-occlusive crises and the steady state condition. Eur Cytokine Netw. 2013;24(1):45–52.

11. Santiago RP, Guarda CC, Figueiredo CVB, et al. Serum haptoglobin and hemopexin levels are depleted in pediatric sickle cell disease patients. Blood Cells Mol Dis. 2018;72:34–36.

12. Noguchi CT, Schechter AN. The intracellular polymerization of sickle hemoglobin and its relevance to sickle cell disease. Blood. 1981;58(6):1057–1068.

13. Hannemann A, Rees DC, Tewari S, Gibson JS. Cation Homeostasis in Red Cells From Patients With Sickle Cell Disease Heterologous for HbS and HbC (HbSC Genotype). EBioMedicine. 2015;2(11):1669–1676.

14. Kuypers FA. Hemoglobin s polymerization and red cell membrane changes. Hematol Oncol Clin North Am. 2014;28(2):155–179.

15. Eaton JW, Skelton TD, Swofford HS, Kolpin CE, Jacob HS. Elevated erythrocyte calcium in sickle cell disease. Nature. 1973;246(5428):105–106.

16. Brugnara C, de Franceschi L, Alper SL. Inhibition of Ca(2+)-dependent K+ transport and cell dehydration in sickle erythrocytes by clotrimazole and other imidazole derivatives. J Clin Invest. 1993;92(1):520–526.

17. Izumo H, Lear S, Williams M, Rosa R, Epstein FH. Sodium-potassium pump, ion fluxes, and cellular dehydration in sickle cell anemia. J Clin Invest. 1987;79(6):1621–1628.

18. Bookchin RM, Lew VL. Effect of a ’sickling pulse’ on calcium and potassium transport in sickle cell trait red cells. J Physiol. 1981;312:265–280.

19. Lew VL, Bookchin RM. Ion transport pathology in the mechanism of sickle cell dehydration. Physiol Rev. 2005;85(1):179–200.

20. Lew VL, Ortiz OE, Bookchin RM. Stochastic nature and red cell population distribution of the sickling-induced Ca2+ permeability. J Clin Invest. 1997;99(11):2727–2735.

21. Milligan C, Rees DC, Ellory JC, et al. A non-electrolyte haemolysis assay for diagnosis and prognosis of sickle cell disease. J Physiol. 2013;591(6):1463–1474.

22. Albuisson J, Murthy SE, Bandell M, et al. Dehydrated hereditary stomatocytosis linked to gain-of-function mutations in mechanically activated PIEZO1 ion channels. Nat Commun. 2013;4:1884.

23. Zarychanski R, Schulz VP, Houston BL, et al. Mutations in the mechanotransduction protein PIEZO1 are associated with hereditary xerocytosis. Blood. 2012;120(9):1908–1915.

24. Cahalan SM, Lukacs V, Ranade SS, Chien S, Bandell M, Patapoutian A. Piezo1 links mechanical forces to red blood cell volume. Elife. 2015;4.

25. Ma S, Cahalan S, LaMonte G, et al. Common PIEZO1 Allele in African Populations Causes RBC Dehydration and Attenuates Plasmodium Infection. Cell. 2018;173(2):443–455.e412.

26. Andolfo I, Alper SL, De Franceschi L, et al. Multiple clinical forms of dehydrated hereditary stomatocytosis arise from mutations in PIEZO1. Blood. 2013;121(19):3925–3935, s3921-3912.

27. Archer NM, Shmukler BE, Andolfo I, et al. Hereditary xerocytosis revisited. Am J Hematol. 2014;89(12):1142–1146.

28. Shmukler BE, Vandorpe DH, Rivera A, Auerbach M, Brugnara C, Alper SL. Dehydrated stomatocytic anemia due to the heterozygous mutation R2456H in the mechanosensitive cation channel PIEZO1: a case report. Blood Cells Mol Dis. 2014;52(1):53–54.

29. Cameron BF, Smariga P. Calcium exchange and calcium-related effects in normal and sickle cell anemia erythrocytes. Prog Clin Biol Res. 1978;20:105–122.

30. Erickson BN, Williams HH, Hummel FC, Lee P, Macy IG. THE LIPID AND MINERAL DISTRIBUTION OF THE SERUM AND ERYTHROCYTES IN THE HEMOLYTIC AND HYPOCHROMIC ANEMIAS OF CHILDHOOD. Journal of Biological Chemistry. 1937;118(3):569–598.

31. Connor WE, Lin DS, Thomas G, Ey F, DeLoughery T, Zhu N. Abnormal phospholipid molecular species of erythrocytes in sickle cell anemia. J Lipid Res. 1997;38(12):2516–2528.

32. Ren H, Okpala I, Ghebremeskel K, Ugochukwu CC, Ibegbulam O, Crawford M. Blood mononuclear cells and platelets have abnormal fatty acid composition in homozygous sickle cell disease. Ann Hematol. 2005;84(9):578–583.

33. Zorca S, Freeman L, Hildesheim M, et al. Lipid levels in sickle-cell disease associated with haemolytic severity, vascular dysfunction and pulmonary hypertension. Br J Haematol. 2010;149(3):436–445.

34. VanderJagt DJ, Trujillo MR, Bode-Thomas F, Huang YS, Chuang LT, Glew RH. Phase angle correlates with n-3 fatty acids and cholesterol in red cells of Nigerian children with sickle cell disease. Lipids Health Dis. 2003;2:2.

35. Romero LO, Massey AE, Mata-Daboin AD, et al. Dietary fatty acids fine-tune Piezo1 mechanical response. Nat Commun. 2019;10(1):1200.

36. Ma S, Dubin AE, Romero LO, et al. Excessive mechanotransduction in sensory neurons causes joint contractures. Science. 2023;379(6628):201–206.

37. Romero LO, Caires R, Kaitlyn Victor A, et al. Linoleic acid improves PIEZO2 dysfunction in a mouse model of Angelman Syndrome. Nat Commun. 2023;14(1):1167.

38. Wu LC, Sun CW, Ryan TM, Pawlik KM, Ren J, Townes TM. Correction of sickle cell disease by homologous recombination in embryonic stem cells. Blood. 2006;108(4):1183–1188.

39. Chien S, Usami S, Bertles JF. Abnormal rheology of oxygenated blood in sickle cell anemia. J Clin Invest. 1970;49(4):623–634.

40. Serjeant GR, Serjeant BE, Milner PF. The irreversibly sickled cell; a determinant of haemolysis in sickle cell anaemia. Br J Haematol. 1969;17(6):527–533.

41. Syeda R, Xu J, Dubin AE, et al. Chemical activation of the mechanotransduction channel Piezo1. Elife. 2015;4.

42. Bae C, Sachs F, Gottlieb PA. The mechanosensitive ion channel Piezo1 is inhibited by the peptide GsMTx4. Biochemistry. 2011;50(29):6295–6300.

43. Bae C, Gnanasambandam R, Nicolai C, Sachs F, Gottlieb PA. Xerocytosis is caused by mutations that alter the kinetics of the mechanosensitive channel PIEZO1. Proc Natl Acad Sci U S A. 2013;110(12):E1162–1168.

44. Romero LO, Caires R, Nickolls AR, Chesler AT, Cordero-Morales JF, Vásquez V. A dietary fatty acid counteracts neuronal mechanical sensitization. Nat Commun. 2020;11(1):2997.

45. Caires R, Garrud TAC, Romero LO, et al. Genetic- and diet-induced ω-3 fatty acid enrichment enhances TRPV4-mediated vasodilation in mice. Cell Rep. 2022;40(10):111306.

46. Yalcinkaya A, Unal S, Oztas Y. Altered HDL particle in sickle cell disease: decreased cholesterol content is associated with hemolysis, whereas decreased Apolipoprotein A1 is linked to inflammation. Lipids Health Dis. 2019;18(1):225.

47. Antwi-Boasiako C, Kusi-Mensah YA, Hayfron-Benjamin C, et al. Total Serum Magnesium Levels and Calcium-To-Magnesium Ratio in Sickle Cell Disease. Medicina (Kaunas). 2019;55(9).

48. Vaisey G, Banerjee P, North AJ, Haselwandter CA, MacKinnon R. Piezo1 as a force-through-membrane sensor in red blood cells. Elife. 2022;11.

49. Nader E, Conran N, Leonardo FC, et al. Piezo1 activation augments sickling propensity and the adhesive properties of sickle red blood cells in a calcium-dependent manner. Br J Haematol. 2023;202(3):657–668.

50. Lewis AH, Grandl J. Mechanical sensitivity of Piezo1 ion channels can be tuned by cellular membrane tension. Elife. 2015;4.

51. Lin YC, Guo YR, Miyagi A, Levring J, MacKinnon R, Scheuring S. Force-induced conformational changes in PIEZO1. Nature. 2019;573(7773):230–234.

52. Yang S, Miao X, Arnold S, et al. Membrane curvature governs the distribution of Piezo1 in live cells. Nat Commun. 2022;13(1):7467.

53. Wu H, Bogdanov M, Zhang Y, et al. Hypoxia-mediated impaired erythrocyte Lands’ Cycle is pathogenic for sickle cell disease. Sci Rep. 2016;6:29637.

54. Kuypers FA, de Jong K. The role of phosphatidylserine in recognition and removal of erythrocytes. Cell Mol Biol (Noisy-le-grand). 2004;50(2):147–158.

55. Awojoodu AO, Keegan PM, Lane AR, et al. Acid sphingomyelinase is activated in sickle cell erythrocytes and contributes to inflammatory microparticle generation in SCD. Blood. 2014;124(12):1941–1950.

56. Tsuchiya M, Hara Y, Okuda M, et al. Cell surface flip-flop of phosphatidylserine is critical for PIEZO1-mediated myotube formation. Nat Commun. 2018;9(1):2049.

57. Shi J, Hyman AJ, De Vecchis D, et al. Sphingomyelinase Disables Inactivation in Endogenous PIEZO1 Channels. Cell Rep. 2020;33(1):108225.

58. Endo J, Arita M. Cardioprotective mechanism of omega-3 polyunsaturated fatty acids. J Cardiol. 2016;67(1):22–27.

59. Wiest EF, Walsh-Wilcox MT, Rothe M, Schunck WH, Walker MK. Dietary Omega-3 Polyunsaturated Fatty Acids Prevent Vascular Dysfunction and Attenuate Cytochrome P4501A1 Expression by 2,3,7,8-Tetrachlorodibenzo-P-Dioxin. Toxicol Sci. 2016;154(1):43–54.

60. Ataga KI, Reid M, Ballas SK, et al. Improvements in haemolysis and indicators of erythrocyte survival do not correlate with acute vaso-occlusive crises in patients with sickle cell disease: a phase III randomized, placebo-controlled, double-blind study of the Gardos channel blocker senicapoc (ICA-17043). Br J Haematol. 2011;153(1):92–104.

61. Ataga KI, Smith WR, De Castro LM, et al. Efficacy and safety of the Gardos channel blocker, senicapoc (ICA-17043), in patients with sickle cell anemia. Blood. 2008;111(8):3991–3997.

62. Daak AA, Dampier CD, Fuh B, et al. Double-blind, randomized, multicenter phase 2 study of SC411 in children with sickle cell disease (SCOT trial). Blood Adv. 2018;2(15):1969–1979.

63. Daak AA, Ghebremeskel K, Hassan Z, et al. Effect of omega-3 (n-3) fatty acid supplementation in patients with sickle cell anemia: randomized, double-blind, placebo-controlled trial. Am J Clin Nutr. 2013;97(1):37–44.

64. Daak AA, Lopez-Toledano MA, Heeney MM. Biochemical and therapeutic effects of Omega-3 fatty acids in sickle cell disease. Complement Ther Med. 2020;52:102482.

65. Tomer A, Kasey S, Connor WE, Clark S, Harker LA, Eckman JR. Reduction of pain episodes and prothrombotic activity in sickle cell disease by dietary n-3 fatty acids. Thromb Haemost. 2001;85(6):966–974.

66. Ridone P, Pandzic E, Vassalli M, et al. Disruption of membrane cholesterol organization impairs the activity of PIEZO1 channel clusters. J Gen Physiol. 2020;152(8).

67. Bhatt DL, Steg PG, Miller M, et al. Cardiovascular Risk Reduction with Icosapent Ethyl for Hypertriglyceridemia. N Engl J Med. 2019;380(1):11–22.

68. Majithia A, Bhatt DL, Friedman AN, et al. Benefits of Icosapent Ethyl Across the Range of Kidney Function in Patients With Established Cardiovascular Disease or Diabetes: REDUCE-IT RENAL. Circulation. 2021;144(22):1750–1759.

69. Harris WS. Understanding why REDUCE-IT was positive - Mechanistic overview of eicosapentaenoic acid. Prog Cardiovasc Dis. 2019;62(5):401–405.

