## Supplementary material for "Enhanced PIEZO1 Function Contributes to the Pathogenesis of Sickle Cell Disease": Table I

**Table 1.** Fatty acid comparison between blood samples of SCD mice fed with either a high-fat or an ω-3 enriched diet.

| | **High-fat diet** | **ω-3 diet** | |
|---|---|---|---|
| **Fatty acids** | µg fatty acid/mg erythrocytes Mean ± SD | µg fatty acid/mg erythrocytes Mean ± SD | *p-value* |
| Dodecanoic acid | 0.95 ± 0.53 | 1.40 ± 1.13 | 0.3965 [1] |
| Myristic acid | 12.54 ± 6.86 | 16.55 ± 7.86 | 0.3681 [1] |
| Pentadecanoic acid | 3.02 ± 1.45 | 3.32 ± 1.67 | 0.7559 [1] |
| Palmitoleic acid | 3.70 ± 2.55 | 102.68 ± 223.61 | 0.0087 [2] |
| Sapienic acid | 3.53 ± 2.39 | 96.00 ± 209.52 | 0.0087 [2] |
| Heptadecenoic acid | 0.63 ± 0.58 | 10.74 ± 23.64 | 0.2616 [2] |
| Heptadecanoic acid | 3.07 ± 1.51 | 3.58 ± 1.85 | 0.6139 [1] |
| Octadecatetraenoic acid | 0.09 ± 0.18 | 0.14 ± 0.083 | 0.0647 [2] |
| Linolenic acid (alpha & gamma) | 0.16 ± 0.12 | 0.18 ± 0.097 | 0.717 [1] |
| Linoleic acid | 17.40 ± 11.31 | 22.26 ± 11.35 | 0.4749 [1] |
| Eicosapentaenoic acid | 0.28 ± 0.30 | 6.26 ± 2.50 | 0.0002 [1] |
| Arachidonic acid | 189.85 ± 423.52 | 9.53 ± 4.42 | 0.3095 [2] |
| Dihomo-γ-linolenic acid | 0.13 ± 0.24 | 0 ± 0 | - |
| Mead acid | 0.26 ± 0.21 | 0.09 ± 0.035 | 0.1293 [1] |
| Eicosadienoic acid | 0.57 ± 0.47 | 0.44 ± 0.25 | 0.9361 [2] |
| Eicosenoic acid | 0.76 ± 0.55 | 8.88 ± 19.33 | 0.2403 [2] |
| Arachidic acid | 1.30 ± 0.74 | 1.50 ± 0.74 | 0.6577 [1] |
| Docosahexaenoic acid | 5.53 ± 4.03 | 9.73 ± 4.62 | 0.0649 [2] |
| Docosapentaenoic acid (ω -3) | 0.58 ± 0.40 | 0.76 ± 0.44 | 0.4732 [1] |
| Docosapentaenoic acid (ω -6) | 0.94 ± 0.53 | 1.29 ± 0.87 | 0.8182 [2] |
| Docosatetraenoic acid | 7.84 ± 17.73 | 0.12 ± 0.069 | 0.0082 [2] |
| Docosatrienoic acid | 2.29 ± 0.96 | 2.38 ± 3.93 | 0.1320 [2] |
| Docosadienoic acid | 0.14 ± 0.12 | 0.18 ± 0.11 | 0.5577 [1] |
| Docosenoic acid | 4.53 ± 3.56 | 61.14 ± 132.56 | 0.1320 [2] |
| Docosanoic acid | 0.89 ± 0.50 | 1.49 ± 0.80 | 0.1538 [1] |
| Nervonic acid | 0.14 ± 0.12 | 0.20 ± 0.077 | 0.3576 [1] |
| Lignoceric acid | 1.16 ± 0.64 | 1.35 ± 0.88 | 0.6842 [1] |
| Hexacosenoic acid | 0.02 ± 0.012 | 0.02 ± 0.023 | 0.9999 [2] |
| Hexacosanoic acid | 0.44 ± 0.28 | 0.36 ± 0.17 | 0.6495 [1] |

[1] Two-tailed Unpaired *t*-test and [2] Two-tailed Mann-Whitney test.
