## Supplementary Figures for "Enhanced PIEZO1 Function Contributes to the Pathogenesis of Sickle Cell Disease"

### Supplementary Material

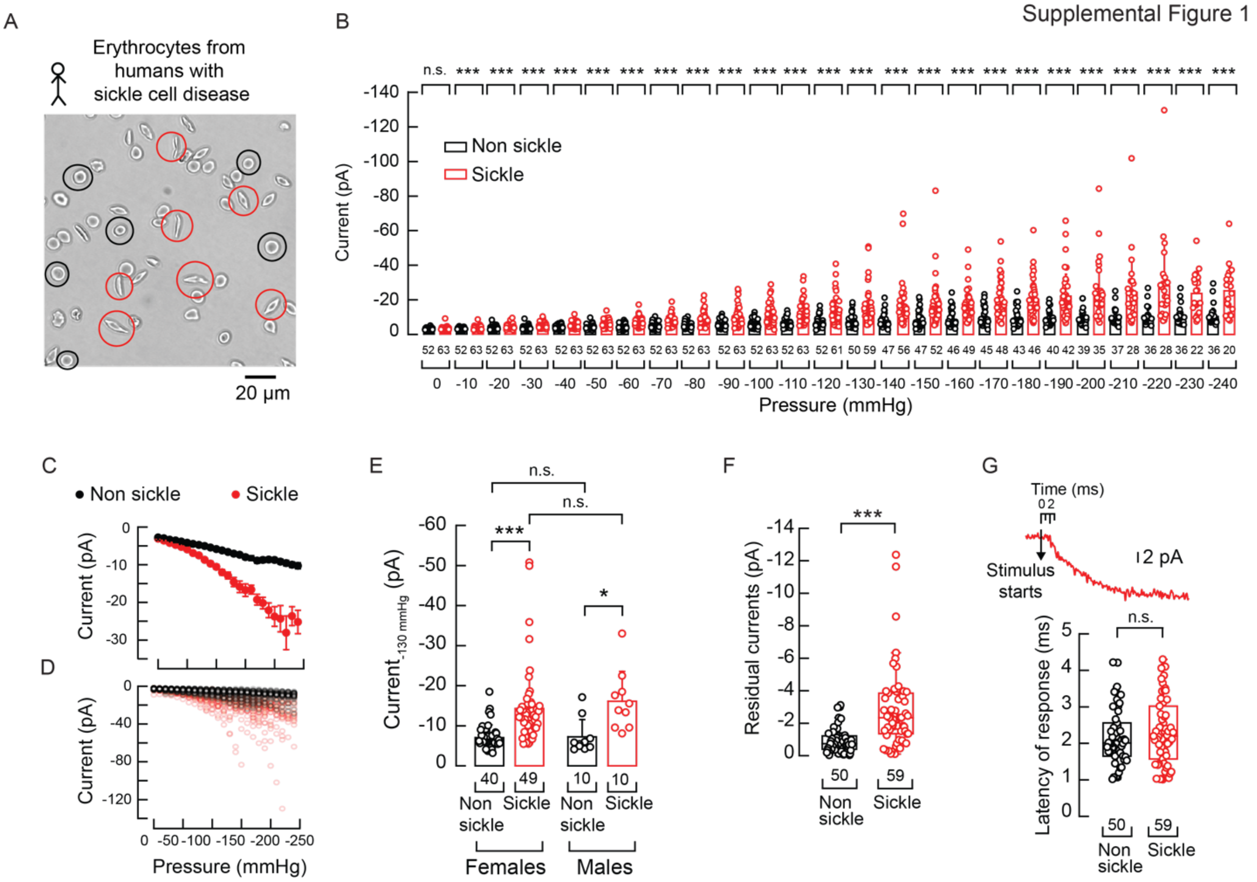


**Supplemental Figure 1. Electrophysiological comparison of non-sickle and sickle erythrocytes from SCD donors. A.** Micrograph of non-sickle and sickle erythrocytes from a fresh blood sample of a donor with sickle cell disease. Encircled in red and black are examples of non-sickle and sickle erythrocytes. Representative of 12 independent donors. **B**. Currents elicited by negative pressure square pulses at a constant voltage of −60 mV. Two-tailed Mann-Whitney test for each pressure step. **C**. Current-pressure relationships (elicited at -60 mV) from sickle (n = 63) and non-sickle (n = 52) erythrocytes, from 12 independent donors. Symbols are mean ± SEM. **D.** Superimposed data points of c. **E**. Mechanocurrents elicited by -130 mmHg from erythrocytes from male and female individuals with sickle cell disease. Two-way ANOVA with Bonferroni multiple comparison test ($F = 0.32$; $p = 0.57$). Bars are mean ± SD. **F**. Current left 30 ms after the -130-mmHg stimulus ends. Two-tailed Mann-Whitney test ($U = 2371$, $p = 1.05^{-8}$). **G**. Top: Representative segment of mechanocurrents elicited by -130 mmHg (at -60 mV) from a sickle human erythrocyte showing the start of the stimulus and the current onset. Bottom: Latency of response to -130 mmHg from. Two-tailed Mann-Whitney test ($U = 1352$, $p = 0.46$). Bars are mean ± SD. Boxplots show the mean, median, and 75$^{th}$ to 25$^{th}$ percentiles. n is denoted above the *x*-axis. Asterisks indicate values significantly different from the control (∗$p < 0.05$, ∗∗$p < 0.01$, and ∗∗∗$p < 0.001$) and n.s. indicates not significantly different.

Supplemental Figure 2

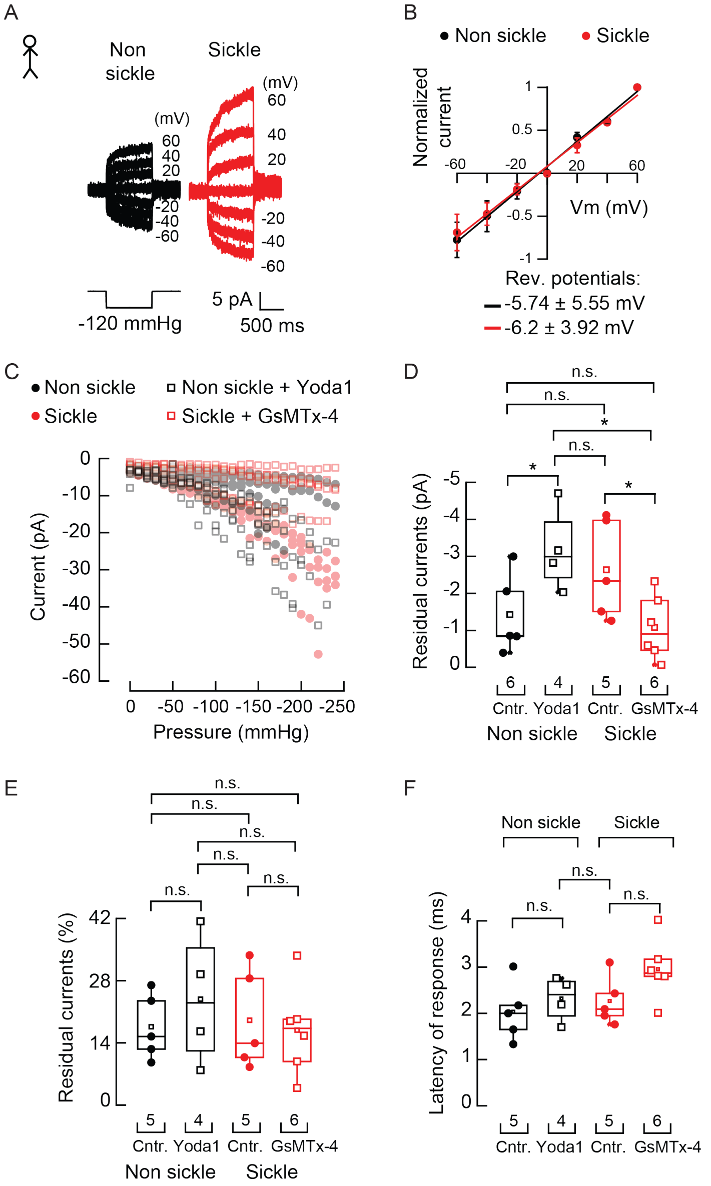


**Supplemental Figure 2. Yoda1 and GsMTx-4 sensitivity of mechanocurrents in non-sickle and sickle erythrocytes from SCD donors. A**. Representative inside-out patch-clamp recordings of mechanically activated currents from non-sickle and sickle erythrocytes. Channel openings were elicited by application of a −120-mmHg-square pulse (bottom) at constant voltages ranging from −60 to +60 mV. **B.** Current-voltage relationships of the currents elicited by -120 mmHg at various membrane potentials from non-sickle (n = 3; rev. potential = -5.74 ± 5.55 mV) and sickle (n = 4; rev. potential = -6.2 ± 3.92 mV) erythrocytes. Symbols are mean ± SD. **C.** Superimposed data points of current-pressure relationships (elicited at -60 mV) from control (DMSO) or Yoda1 (30 µM)-exposed non-sickle and control or GsMTx-4 (7 µM)-exposed sickle erythrocytes. **D.** Current left 30 ms after the -130-mmHg stimulus ends from control (DMSO) or Yoda1 (30 µM)-exposed non-sickle and control or GsMTx-4 (7 µM)-exposed sickle erythrocytes. Kruskal-Wallis ($H = 8.26$, $p = 0.041$) with Dunn's multiple comparison test. **E.** % of peak current 30 ms after the -130-mmHg stimulus ends from control (DMSO) or Yoda1 (30 µM)-exposed non-sickle and control or

GsMTx-4 (7 μM)-exposed sickle erythrocytes. Kruskal-Wallis (H = 0.6048, $p$ = 0.8953) with Dunn's multiple comparison test **F.** Latency of response to -130 mmHg of control (DMSO) or Yoda1-exposed non-sickle and control or GsMTx-4 exposed sickle human erythrocytes. Kruskal-Wallis (H = 5.84, $p$ = 0.12) with Dunn's multiple comparison test. Boxplots show the mean, median, and 75$^{th}$ to 25$^{th}$ percentiles. n is denoted above the $x$-axis. n.s. indicates not significantly different.

Supplemental Figure 3

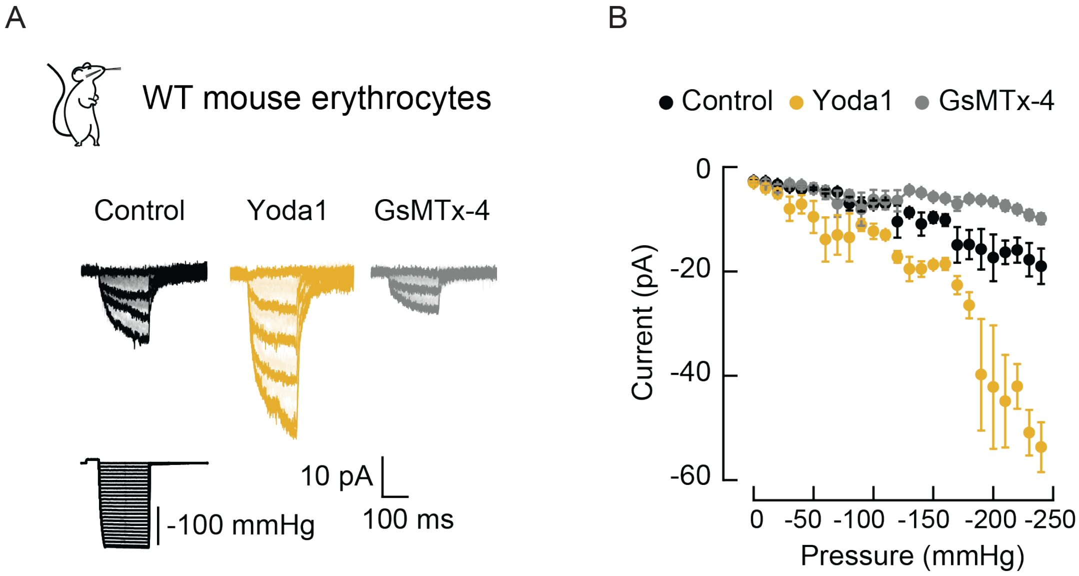


**Supplemental Figure 3. Mouse erythrocytes are Yoda1 and GsmTx-4 sensitive. A**. Representative inside-out recordings elicited by negative pressure square pulses at a constant voltage of −60 mV from wild-type (WT) mouse erythrocytes challenged with DMSO (control), Yoda1 (30 μM), or GsMTx-4 (7 μM). **B**. Current-pressure relationships (elicited at -60 mV) from wild-type (WT) mouse erythrocytes challenged with DMSO (control; n = 10), Yoda1 (n = 8), or GsMTx-4 (n = 6). Symbols are mean ± SEM.

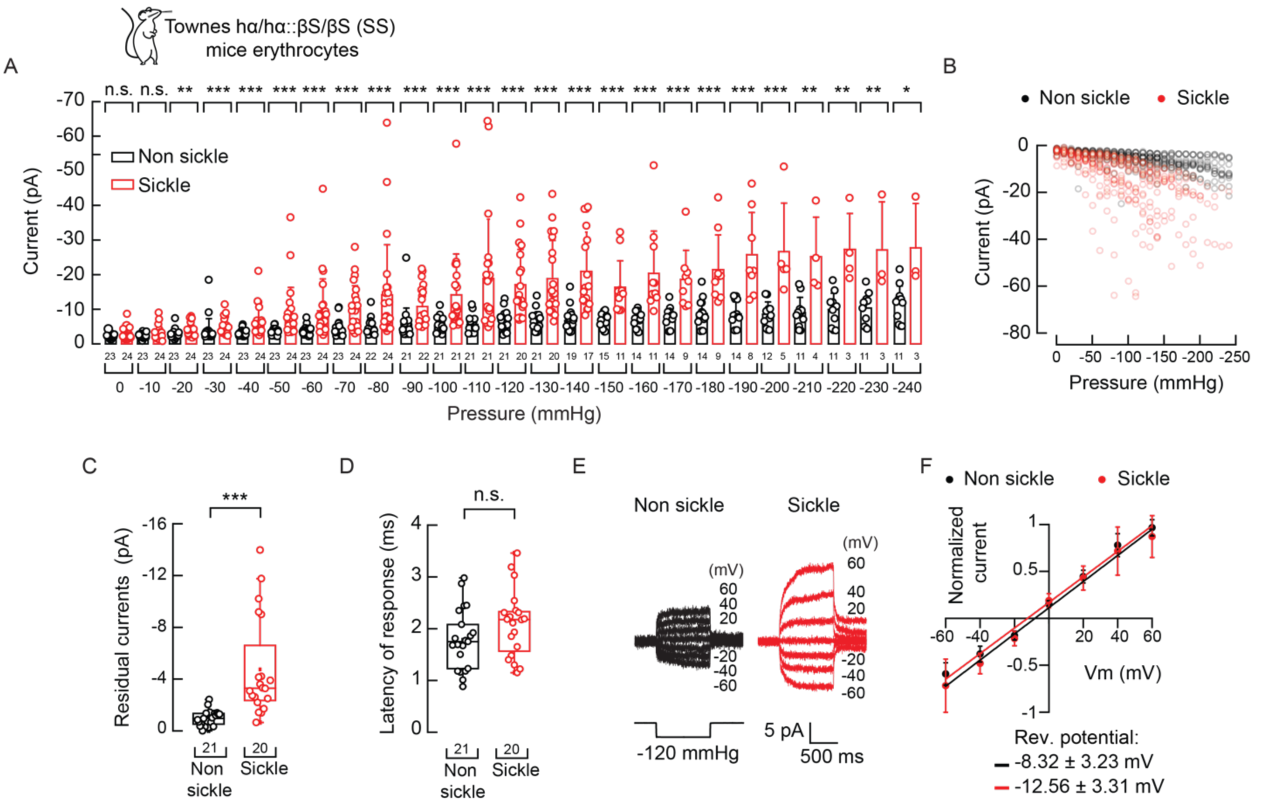


**Supplemental Figure 4. Electrophysiological analysis of non-sickle and sickle erythrocytes from Townes SCD mice. A.** Current elicited by negative pressure square pulses at a constant voltage of −60 mV from non-sickle and sickle erythrocytes. Two-tailed Mann-Whitney test for each pressure step. **B.** Superimposed data points of the current-pressure relationships (elicited at -60 mV) from non-sickle (n = 24) and sickle (n = 23) erythrocytes from 7 mice, shown in Figure 3C. Symbols are mean ± SEM. **C.** Current left 30 ms after the -130-mmHg stimulus ends. Two-tailed Mann-Whitney-test (U = 24, $p$ = $5.44^{-8}$). **D.** Latency of response to -130 mmHg. Two-tailed Unpaired $t$-test ($t$ = 1.67, $p$ = 0.104). **E.** Representative inside-out patch-clamp recordings of mechanically activated currents from non-sickle (n = 3) and sickle erythrocytes (n = 6). Channel openings were elicited by application of a −120-mmHg-square pulse (bottom) at constant voltages ranging from −60 to +60 mV. **F.** Current-voltage relationships of the currents elicited by -120 mmHg at various membrane potentials from non-sickle (n = 3; rev. potential = -8.32 ± 3.23 mV) and sickle (n = 6; rev. potential = -12.56 ± 3.31 mV) erythrocytes Symbols are mean ± SD. Bars are mean ± SD. Boxplots show the mean, median, and $75^{th}$ to $25^{th}$ percentiles. n is denoted above the $x$-axis. Asterisks indicate values significantly different from the control (∗$p < 0.05$, ∗∗$p < 0.01$ and ∗∗∗$p < 0.001$) and n.s. indicates not significantly different.

Supplemental Figure 5

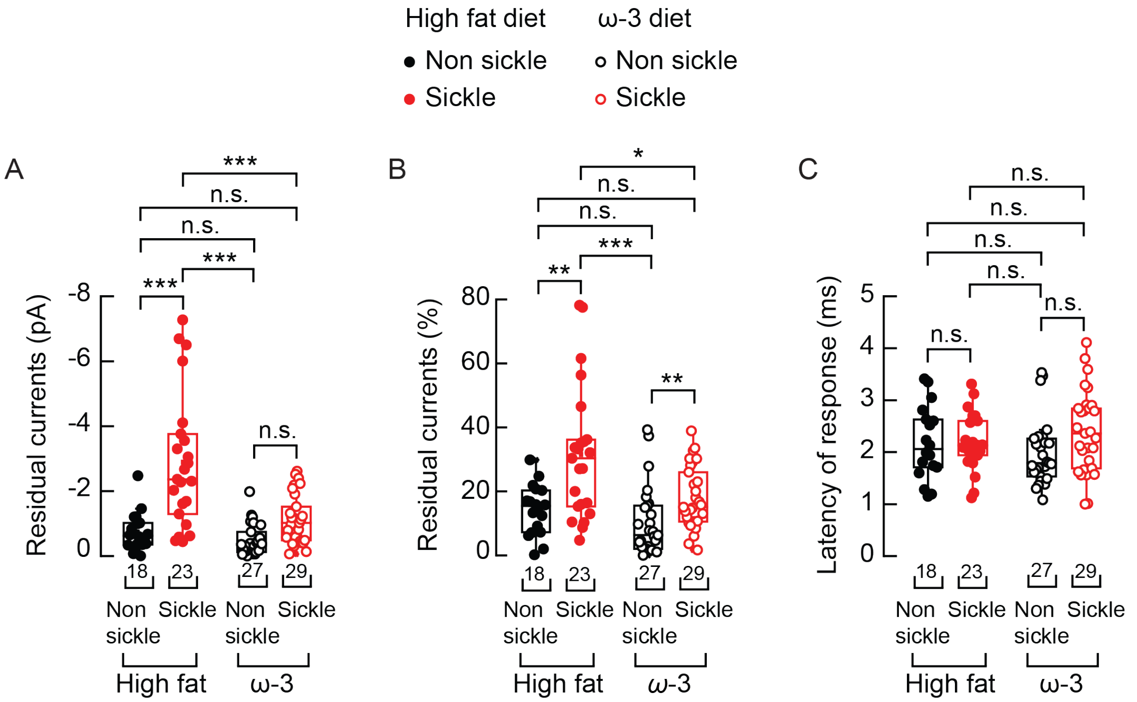


**Supplemental Figure 5. Electrophysiological analysis of non-sickle and sickle erythrocytes post-dietary intervention. A.** Current left 30 ms after the -130-mmHg stimulus ends from non-sickle and sickle erythrocytes of mice fed with isocaloric high-fat or ω-3 enriched diets. Two-way ANOVA with Tukey multiple comparison test ($F = 16.003$; $p = 1.27^{-4}$). **B.** Percentage of peak current 30 ms after the -130-mmHg stimulus ends from non-sickle and sickle erythrocytes of mice fed with isocaloric high-fat or ω-3 enriched diets. Kruskal-Wallis ($H = 23.98$, $p = 2.52^{-5}$) with Dunn's multiple comparison test. **C.** Latency of response to -130 mmHg from non-sickle and sickle erythrocytes of mice fed with isocaloric high-fat or ω-3 enriched diets. Two-way ANOVA with Tukey multiple comparison test ($F = 1.77$; $p = 0.19$). Boxplots show the mean, median, and 75$^{th}$ to 25$^{th}$ percentiles. n is denoted above the *x*-axis. Asterisks indicate values significantly different from the control ($*p < 0.05$, and $***p < 0.001$) and n.s. indicates not significantly different.

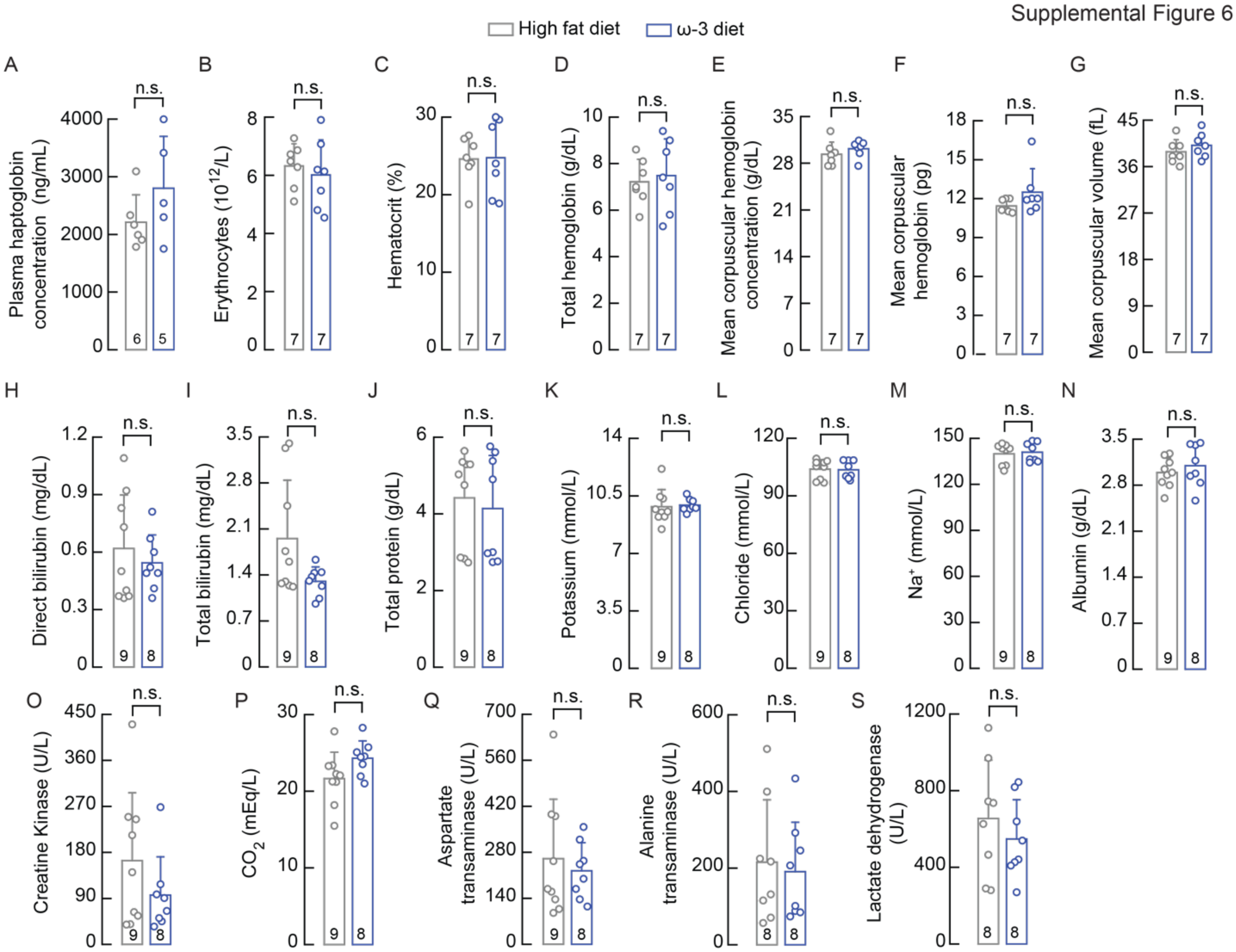


**Supplemental Figure 6. Hematology analyses of the Townes mouse model of sickle cell disease fed with isocaloric high-fat or ω-3 enriched diets. A.** Plasma haptoglobin concentration. Two-tailed Unpaired *t*-test ($t$ = 1.3989, $p$ = 0.1953). **B.** Erythrocytes concentration. Two-tailed Unpaired *t*-test ($t$ = 0.5714, $p$ = 0.5783). **C.** Hematocrit percentage. Two-tailed Unpaired *t*-test ($t$ = 0.0648, $p$ = 0.9494). **D.** Total hemoglobin concentration. Two-tailed Unpaired *t*-test ($t$ = 0.3883, $p$ = 0.7046). **E.** Mean corpuscular hemoglobin concentration. Two-tailed Unpaired *t*-test ($t$ = 0.9714, $p$ = 0.3505). **F.** Mean corpuscular hemoglobin. Two-tailed Mann-Whitney test (U = 13, $p$ = 0.1539). **G.** Mean corpuscular volume. Two-tailed Unpaired *t*-test ($t$ = 1.013, $p$ = 0.3312). **H.** Direct bilirubin concentration. Two-tailed Unpaired *t*-test ($t$ = 0.6823, $p$ = 0.5055). **I.** Total bilirubin concentration. Two-tailed Unpaired *t*-test with Welch correction ($t$ = 2.123, $p$ = 0.0628). **J.** Total protein concentration. Two-tailed Mann-Whitney test (U = 35, $p$ = 0.9616). **K.** Serum potassium concentration. Two-tailed Unpaired *t*-test ($t$ = 0.2379, $p$ = 0.8152). **L.** Serum chloride concentration. Two-tailed Unpaired *t*-test ($t$ = 0.1128, $p$ = 0.9117). **M.** Serum sodium concentration. Two-tailed Mann-Whitney test (U = 28, $p$ = 0.4807). **N.** Serum albumin concentration. Two-tailed Unpaired *t*-test ($t$ = 0.7903, $p$ = 0.4416). **O.** Serum creatine kinase (CK) concentration Two-tailed Unpaired *t*-test ($t$ = 1.269, $p$ = 0.2237). **P.** Serum $CO_2$ concentration. Two-tailed Unpaired *t*-test ($t$ = 1.861, $p$ = 0.0824). **Q.** Serum aspartate transaminase concentration. Two-tailed Unpaired *t*-test ($t$ = 0.5196, $p$ = 0.6109). **R.** Serum alanine transaminase concentration. Two-tailed Unpaired *t*-test ($t$ = 0.3413, $p$ = 0.7379). **S.** Serum lactate dehydrogenase. Two-tailed Unpaired *t*-test ($t$ = 0.8229, $p$ = 0.4244). Bars are mean ± SD. n is denoted above the $x$-axis. n.s. indicates not significantly different.

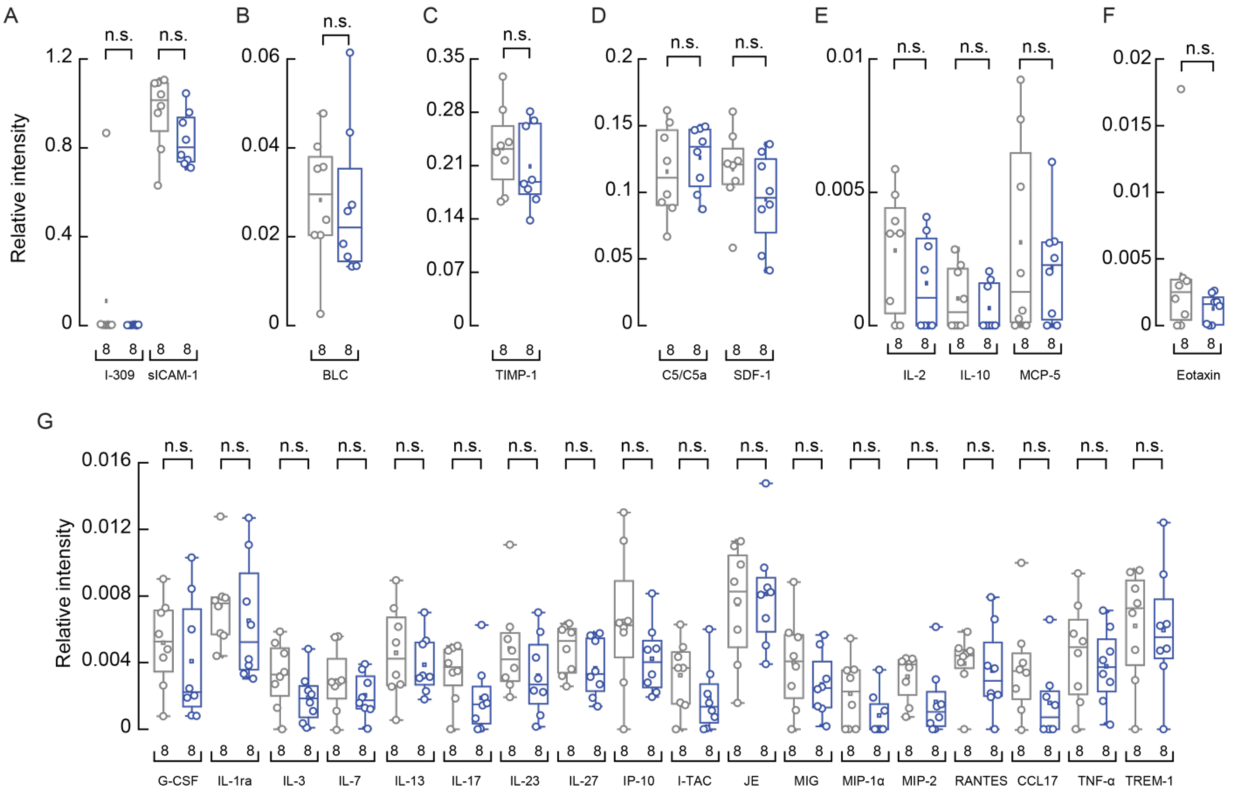


**Supplemental Figure 7. Cytokine profile analyses of the Townes mouse model of sickle cell disease fed with isocaloric high-fat or ω-3 enriched diets. A.** Relative intensity of serum I-309 and soluble intercellular adhesion molecule (sICAM). Two-tailed Mann-Whitney test (I-309: U = 15 and $p = 0.083$) and Two-tailed Unpaired *t*-test (sICAM: $t = 1.681$ and $p = 0.115$). **B.** Relative intensity of serum B lymphocyte chemoattractant (BLC). Two-tailed Unpaired *t*-test ($t = 0.1228$ and $p = 0.9040$). **C.** Relative intensity of serum tissue inhibitor of metalloproteinases-1 (TIMP-1). Two-tailed Unpaired *t*-test ($t$ = 0. 0.8707 and $p$ = 0.3986). **D.** Relative intensity of serum C5/C5a and stromal cell-derived factor-1 (SDF-1). Two-tailed Unpaired *t*-test (C5/C5a: $t = 0.71$ and $p = 0.4894$; SDF-1: $t = 1.39$ and $p = 0.1862$). **E**. Relative intensity of serum interleukin-2 (IL-2), interleukin-10 (IL-10) and monocyte chemoattractant-5 (MCP-5). Two-tailed Mann-Whitney test (IL-2: U = 21 and $p = 0.2576$, IL-10: U = 26 and $p = 0.5245$) and Two-tailed Unpaired *t*-test (MCP-5: $t = 0.6242$ and $p = 0.5425$). **F**. Relative intensity of serum eotaxin. Two-tailed Mann-Whitney test (U = 21 and $p = 0.2667$). **G.** The array panel includes relative intensities of multiple cytokines. Two-tailed Unpaired *t*-test (IL-3: $t = 1.557$ and $p = 0.1417$; IL-7: $t = 1.067$ and $p = 0.3041$; IL-13: $t = 0.6071$ and $p = 0.5535$; IL-17 : $t = 1.615$ and $p = 0.1285$; IL-23: $t = 1.231$ and $p = 0.2387$; IL-27: $t = 1.426$ and $p = 0.1756$; IP-10: $t = 1.37$ and $p = 0.1923$; I-TAC: $t = 1.355$ and $p = 0.1969$; JE: $t = 0.3524$ and $p = 0.7298$; MIG: $t = 1.0918$ and $p = 0.2908$; MIP-2: $t = 1.712$ and $p = 0.109$; RANTES: $t = 0.3872$ and $p = 0.7044$; CCL17: $t = 1.594$ and $p = 0.1334$; TNF-α: $t = 0.5883$ and $p = 0.5657$; TREM-1: $t = 0.1459$ and $p = 0.8861$) and Two-tailed Mann-Whitney test (G-CSF: U = 25 and $p = 0.5054$; IL-1ra: U = 21 and $p = 0.2786$; CCL3/MIP-1α: U = 20.5 and $p = 0.2173$). Boxplots show the mean, median, and 75$^{th}$ to 25$^{th}$ percentiles. n is denoted above the *x*-axis. n.s. indicates not significantly different.
